# Fusiform face area development correlates with development in higher-order social brain regions

**DOI:** 10.64898/2026.03.10.710863

**Authors:** Lorena Jiménez-Sánchez, Melissa Thye, Hilary Richardson

## Abstract

3.

The fusiform face area (FFA) preferentially responds to faces within the first months of life. One hypothesis is that higher-order social responses in middle medial prefrontal cortex (MMPFC) or face responses in superior temporal sulcus (STS) drive the development of face-selective responses in FFA, with right-hemisphere dominance in FFA eventually arising from lateralised connections to these regions. Another hypothesis proposes an innate face template in the amygdala guides attention to face-like shapes. This study opportunistically examined the development of the FFA, MMPFC, STS, and amygdala in childhood using an open cross-sectional movie-viewing fMRI dataset with 3-12-year-olds (N=117, M=6.77 years) and adults (N=33, M=24.77 years). We tested for correlations between FFA development and development in MMPFC, STS, and amygdala on the premise that associations between these regions may be observable even in children, and such associations could constrain hypotheses and analytic approaches in future studies with infants. First, we measured functional maturity-how similar each child’s response to the movie was to an adult average response timecourse. In all regions, older children’s responses were more adult-like. Next, we tested whether FFA maturity correlated with functional connectivity with, or functional maturity of, MMPFC, STS, or amygdala. Children with more mature right FFA responses showed stronger right FFA-right MMPFC connectivity. Children with more mature FFA responses also had more mature STS responses, bilaterally. This study provides preliminary evidence that FFA co-develops with higher-order social brain regions and specific metrics to take forward in future research with infants.

**Highlights:**

- What drives face selective responses in FFA is the subject of recent debate.
- 117 children aged 3 to 12 years watched a short movie while undergoing fMRI.
- Right FFA development correlated with functional connectivity to right MMPFC
- FFA development correlated with STS development, bilaterally.
- FFA codevelops with higher-order social brain regions (controlling for age).

## 5. Introduction

Faces are salient stimuli in our environment and an important source for recognising and learning about other people. In adults, a distributed network of cortical regions respond preferentially to faces (Haxby et al., 2000, Kanwisher et al., 1997), including bilateral superior temporal sulcus (STS), occipital face area, and fusiform face area (FFA); these responses are more dominant in the right hemisphere (Kanwisher et al., 1997, Behrmann and Plaut, 2020). In particular, the right FFA demonstrates the most consistent and robust face-selective activation relative to control stimuli (e.g., bodies, objects, and scenes) in adults (Downing et al., 2006, Grill-Spector et al., 1998, Kanwisher et al., 1997) and functional connectivity between face regions is stronger in the right hemisphere from infancy (Lesinger et al., 2023). The FFA, and other face-selective brain regions, appear to have face-selective responses by two months of age (Kosakowski et al., 2024), with no observable effect of age among 2-9 month old infants. However, there is also evidence for continued increases in the selectivity of FFA responses for faces, primarily via reduced responses to other visual categories, in childhood (Cantlon et al., 2011, Nordt et al., 2021, Peelen et al., 2009, Natu et al., 2016, Gomez et al., 2017).

What are the developmental drivers of face-selective responses in FFA? Powell et al. (2018) articulated three non-mutually exclusive hypotheses derived from research to date. The first hypothesis proposes that infants choose to look at faces to engage in positive, self-relevant social interactions. In this account, responses to positive, self-relevant social stimuli in middle medial prefrontal cortex (MMPFC) drive attention to faces and consequently the development of face-selective FFA responses. MMPFC responds to socially relevant stimuli (e.g., familiar faces or voices, mutual gaze) in adults (Amodio and Frith, 2006, Gobbini and Haxby, 2007, Van Den Bos et al., 2007) and infants (Grossmann et al., 2008, Imafuku et al., 2014, Naoi et al., 2012, Urakawa et al., 2015), and shows selective responses to faces in two-month old infants (Kosakowski et al., 2024). There is some evidence that MMPFC is involved in face processing from birth: an electroencephalography (EEG) study in newborns reported face-specific neural responses that were localised to several cortical regions, including MMPFC (Buiatti et al., 2019). There is also evidence for “top-down” influences from MMPFC and STS to FFA responses in adults asked to switch between reporting the emotional valence and the age of a face (Anzellotti et al., 2017). This hypothesis could plausibly explain right-hemisphere dominance for face processing, which could arise from laterally-biased connectivity with “higher order” regions including MMPFC (Powell et al., 2018), similar to how pre-existing connectivity to higher order language regions explains the left-lateralised development of the visual word form area (Saygin et al., 2016, Op de Beeck et al., 2019). More recent theoretical accounts extend this view and propose that hemispheric asymmetries emerge through an interaction between such neural constraints and experience viewing faces (Behrmann et al., 2025). Infant fMRI studies have yet to detect right-lateralised face responses (Kosakowski et al., 2024), but a right-hemisphere bias in face responses has been observed in school-aged children and adults (Liu et al., 2024, Kanwisher et al., 1997; also see Thome et al., 2022).

The second hypothesis described by Powell et al. (2018) is that the brain contains an innate, subcortical “face template” that directs infants’ attention to face-like shapes (Morton and Johnson, 1991), driving the development of face-selective cortical regions. While there is currently no direct evidence for a subcortical “face template”, studies with adults support the existence of a rapid subcortical route for face processing involving the amygdala (Krolak-Salmon et al., 2004). This route mainly reflects subcortical responses to low spatial frequencies of visual stimuli (Vuilleumier et al., 2003) which may influence cortical processing, including in fusiform gyrus (Morris et al., 1998). However, many of the aforementioned results involve experiments using fearful faces, so it remains unclear if amygdala responses reflect matches between face stimuli and a face template, or if they reflect processing of other aspects of these stimuli, such as fear (Johnson, 2005), emotional valence (Liu et al., 2022), salience and/or biological/social relevance (Adolphs, 2008). Additionally, very little research investigates amygdala responses in human infants, likely because it is not possible to measure subcortical responses using functional near-infrared spectroscopy (fNIRS) or EEG – which are more tolerant to participant motion than fMRI. To our knowledge, only one study has used fMRI to measure amygdala responses to faces in infancy. While fMRI affords the high spatial resolution and full-brain coverage necessary for studying the amygdala, results were inconclusive (Kosakowski et al., 2021). Kosakowski et al. reported difficulties parcellating the amygdala using structural MRI data, given poor tissue contrast in infants, and difficulties coregistering the parcellated amygdala with awake infant fMRI data (given time-resolution trade-offs made during the development of awake infant fMRI sequences); because subcortical responses are smaller in magnitude than cortical responses, the consequences of these methodological challenges are exacerbated. It remains possible that the amygdala responds to faces even in infants, but these responses have not yet been detected.

The third hypothesis described by Powell et al. (2018) suggests that infants’ visual experiences co-activate distinct neural populations with preferences for different low-level visual image statistics (e.g., rectilinearity vs curvilinearity), and repeated co-activation over time drives the development of preferential responses to faces (Livingstone et al., 2017, Arcaro et al., 2017). In particular, parents often interact with their infants face-to-face, so infants acquire extensive experience with face images on their fovea at a close distance. Neurons in the extrastriate cortex with preferences for foveal input, curvilinearity, and low spatial frequency are then repeatedly co-activated over time and, perhaps, become selective for face stimuli in virtue of these characteristics. However, this mechanism predicts bilaterally symmetric face responses, and so this hypothesis cannot yet explain (eventual) right-lateralised face responses.

We conducted opportunistic analyses of naturalistic movie-viewing data with 3–12-year-old children to test predictions of the first two hypotheses and investigate correlates of face-selective responses in right FFA. While our dataset was cross-sectional and cannot reveal causal mechanisms, functional activity and connectivity in childhood could reflect early and historical functioning and connectivity of face-responsive regions. If right FFA specialisation is driven by connectivity to or development of ’top down’ regions such as MMPFC, this causal history might be reflected in a concurrent correlation between FFA development and its functional connectivity to MMPFC in children. This premise is supported by machine learning studies which use resting-state connectivity to predict task-based brain activity in children (Bernstein-Eliav and Tavor, 2024). While evidence from infants suggests early development of face-selective responses (Deen et al., 2017, Kosakowski et al., 2024), there is also evidence for developmental change in cortical face-selectivity in childhood, with some neurotypical children showing no discernible selectivity in FFA by age 5-7 (Cantlon et al., 2011, Cohen et al., 2019, Golarai et al., 2007, Gomez et al., 2017, Scherf et al., 2007). Our approach could therefore plausibly reveal neural correlates of individual differences in developing face-selective responses among children.

In preregistered analyses, we characterised functional responses of bilateral FFA, MMPFC, and amygdala to faces and tested whether functional maturity of FFA correlates with its functional connectivity to MMPFC and amygdala. Primary analyses focused on correlates of right FFA development, given evidence for right-hemisphere dominance for face-selective responses (Kanwisher et al., 1997, Behrmann and Plaut, 2020). In non-preregistered analyses, we conducted similar analyses testing for correlations between STS and FFA development; the STS is another higher-order face-selective region that could plausibly shape development of FFA (Anzellotti et al., 2017, Deen et al., 2017, Kosakowski et al., 2024). In additional non-preregistered analyses, we further examined correlations between the functional maturity of the right FFA and bilateral MMPFC, amygdala, and STS.

## 6. Materials and methods

This study involved opportunistic analyses of an open fMRI dataset downloaded from OpenNeuro (https://openneuro.org/datasets/ds000228; Richardson et al., 2018). Analyses were preregistered on the Open Science Framework (https://osf.io/hytsr). Deviations are reported where relevant and summarised in **Table S1**.

### 6.1 Participants and data

The dataset includes children (N = 122; ages 3.5–12 years), and adults (N=33; ages 18–39 years). Five children were excluded from analyses for preprocessing issues (segmentation error likely due to a priori defacing issues, n=1) or not meeting participant motion exclusion criteria (see section 7, n=4). The final sample included 117 children (M(SD)=6.77(2.33) years of age; 62 females) and 33 adults (M(SD)=24.77(5.31) years of age; 20 females).

All participants were recruited from the local community surrounding Massachusetts Institute of Technology (MIT) (Cambridge, MA, USA), gave written consent (parent/guardian consent and child assent was received for all child participants), and had normal or corrected-to-normal vision. Recruitment and experiment protocols were approved by the Committee on the Use of Humans as Experimental participants (COUHES) at MIT.

Primary analyses were conducted using data from children; data from adults were used only to identify timepoints for response magnitudes to face and scene events and to calculate functional maturity in children (section 7).

### 6.2 fMRI stimuli

All participants watched a silent version of “Partly Cloudy” (Reher and Sohn, 2009), a 5.6-min animated movie. A short description of the plot can be found online (https://www.pixar.com/partlycloudy#partly-cloudy-1). The stimulus was preceded by 10 s of rest and subtended 17.62 x 13.07° of visual angle.

### 6.3 fMRI data acquisition

Data acquisition procedures and parameters have been described previously (Richardson et al., 2018). Briefly, children completed a mock scan to acclimate to the scanner and learn to stay still. Young participants could hold a stuffed animal for comfort and to reduce movement. During scanning, an experimenter stood near the bore to ensure children stayed awake, attended to the movie, and remained still.

Whole-brain structural and functional MRI data were acquired using a 3-Tesla Siemens Tim Trio scanner at the Athinoula A. Martinos Imaging Centre at MIT. Children under age 5 used custom 32-channel phased-array head coils, while older participants used the standard Siemens 32-channel head coil. T1-weighted structural images were collected with 1 mm isotropic voxels. Functional data were obtained in a single run with a gradient-echo EPI sequence sensitive to Blood Oxygen Level Dependent (BOLD) contrast in 32 interleaved near-axial slices aligned with the anterior/posterior commissure, and covering the whole brain (repetition time, TR: 2 s; 168 volumes). Participants were recruited for different studies and had small differences in voxel size and slice gaps, so all functional data were up-sampled in normalised space to 2 mm isotropic voxels. Prospective acquisition correction was used to adjust for head motion one TR back (Thesen et al., 2000).

### 6.4 fMRI data analysis

#### Preprocessing

Data were preprocessed using fMRIPrep 24.0.0 (Esteban et al., 2019, Markiewicz et al., 2024) which is based on Nypipe 1.8.6 (Gorgolewski et al., 2011). Briefly, data were run through the complete fMRIPrep workflow, which includes brain-skull separation, brain tissue segmentation, spatial normalisation, and confound estimation. A detailed description can be found at **Appendix S1**. Data were then smoothed using a Gaussian filter with a 5 mm kernel.

#### Motion treatment

FMRIPrep outputs were visually inspected and subjected to a quality control workflow to exclude data with excessive participant movement prior to first-level modelling. Outlier volumes were defined as those with (1) framewise displacement (FD) > 1 and (2) using the following thresholds within rapidart, the nipype implementation of the ART toolbox: more than 1mm composite motion relative to the previous volume or a fluctuation in global signal exceeding 3 standard deviations from the mean global signal. If one-third or more of the volumes within the functional run were identified as motion outliers, participants were excluded from analyses (n=4 children). Although preregistered, the threshold of standardised DVARS > 1.5 was not applied, as it was not an analysis-informed threshold and resulted in the exclusion of participants with otherwise acceptable data quality (https://osf.io/hytsr); these participants have been included in prior publications using this dataset (Richardson et al., 2018, Yates et al., 2021). Mean FD was uncorrelated with age among children (Spearman correlation test: rs(115)=-0.06, p=0.510) and was included as a covariate in statistical analyses to account for participant motion (see section 9).

#### Region of interest (ROI) definition

We analysed responses in subject-specific left and right FFA, MMPFC, amygdala and STS ROIs (8 ROIs total). Analyses of STS were conducted after our preregistered analyses with other ROIs were completed; we did not preregister analyses of STS responses. FFA, MMPFC and STS ROIs were defined functionally using a procedure described previously (Kamps, Richardson et al., 2022). In this approach, the average response timecourse from localiser-defined functional ROIs (fROIs) in independent adult samples (‘reference timecourses’) are regressed on movie timecourses for each child. This provides an estimate of how similar a voxel’s timecourse is to the adult reference timecourse. Subject-specific functional ROIs are then defined as the highest-ranking voxels to contrasts of previously extracted, localiser-defined fROI timecourses chosen to reflect the domain-selectivity of each region. Relative to approaches that use particular movie scenes as events (e.g., face events, scene events) to define contrasts, this approach leverages the full response timecourse for ROI definition. Here, left and right FFA ROIs (defined with faces > objects in the original localiser task with adults) were defined as the top 80 voxels to the contrast of (left or right) FFA > (left or right) lateral occipital cortex (LOC) timecourses, within large left and right FFA search spaces (Julian et al., 2012). Left and right MMPFC ROIs (defined with false beliefs > false photographs in the original localiser task with adults) were defined as the top 80 voxels to the contrast of (left or right) MMPFC > (left or right) secondary somatosensory cortex (S2), within a large bilateral MMPFC search space (Dufour et al., 2013); https://saxelab.mit.edu/use-our-theory-mind-group-maps/) split by hemisphere (for left and right MMPFC search spaces, see https://osf.io/mxkag/). We targeted middle MPFC (MMPFC) rather than dorsal or ventral MPFC based on reviewing whole-brain maps of responses to faces in individual infants reported previously (e.g., Figure 3 in Kosakowski et al., 2024). Left and right STS ROIs (defined with faces > objects in the original localiser task with adults) were defined as the top 80 voxels to the contrast of (left or right) STS > (left or right) LOC timecourses, within large left and right STS search spaces (Julian et al., 2012). For clarity, further details about the adult reference timecourses used for fROI definition are provided in **Appendix S2.** First-level modelling carried out for fROI definition included rapidart-identified outlier volumes, the top five aCompCor components and adult reference timecourses for all localiser-defined fROIs as additional regressors; all timecourses were included in one model, following previous research (Kamps, Richardson et al., 2022). Data were high-pass filtered (0.01 Hz).

**Figure 1.**
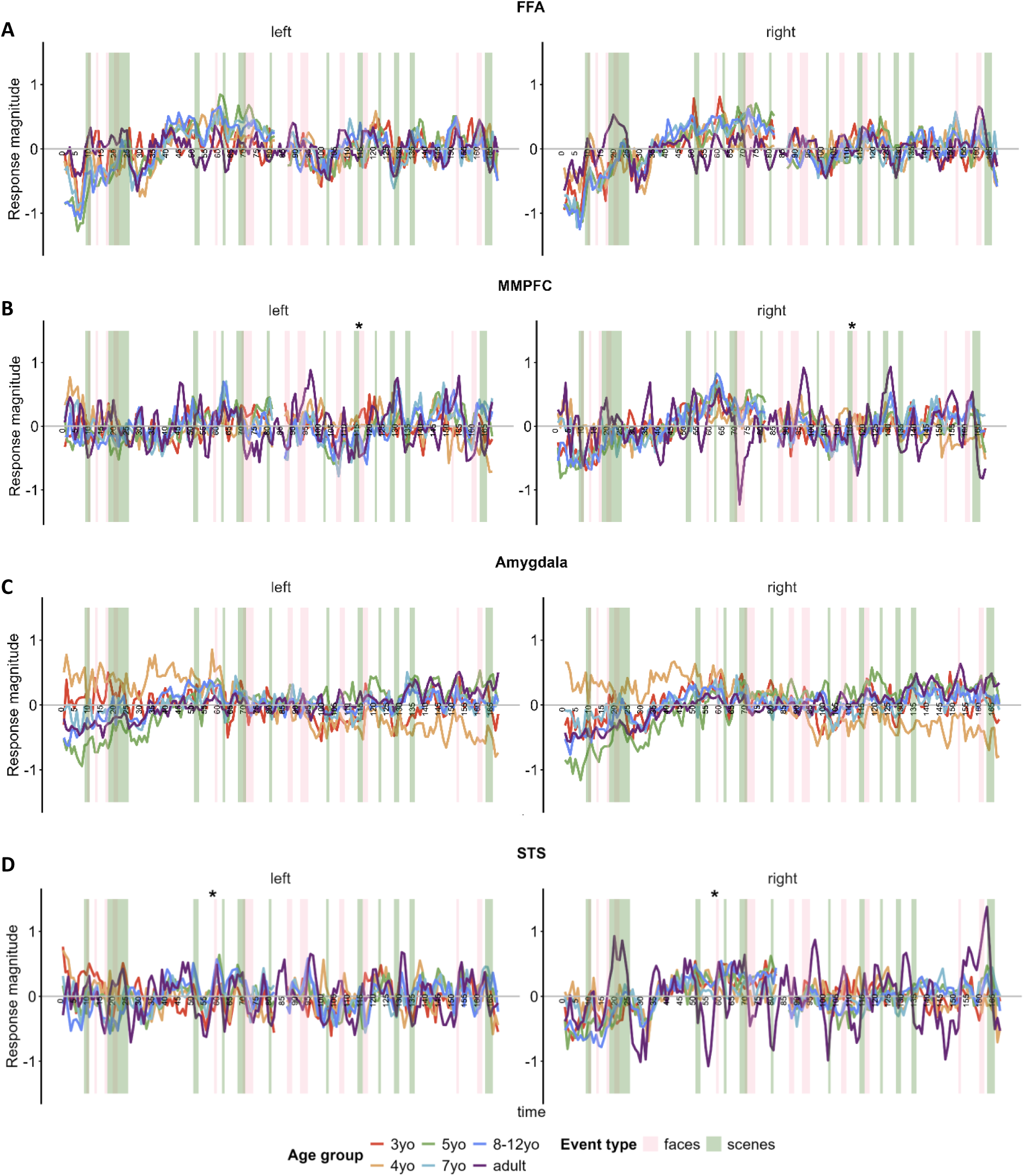
Average timecourse per age group for A) FFA, B) MMPFC, C) Amygdala, and D) STS during viewing of ‘Partly Cloudy’. Each timepoint along the x-axis corresponds to a single TR (2 s). Shaded blocks show timepoints identified as face (pink) and scene (green) events in a reverse correlation analysis conducted on adults in prior research (Kamps, Richardson et al., 2022). Left and right panels show measures for the left and right hemisphere, respectively. An asterisk above the face event F10 and F05 indicate that the MMPFC and STS response magnitude for this event, respectively, showed significant age effects after correcting for multiple comparisons (α=0.004). For all plots, N=117. FFA = Fusiform Face Area; MMPFC = Middle Medial Prefrontal Cortex; STS = Superior Temporal Sulcus; yo = year old.

**Figure 2.**
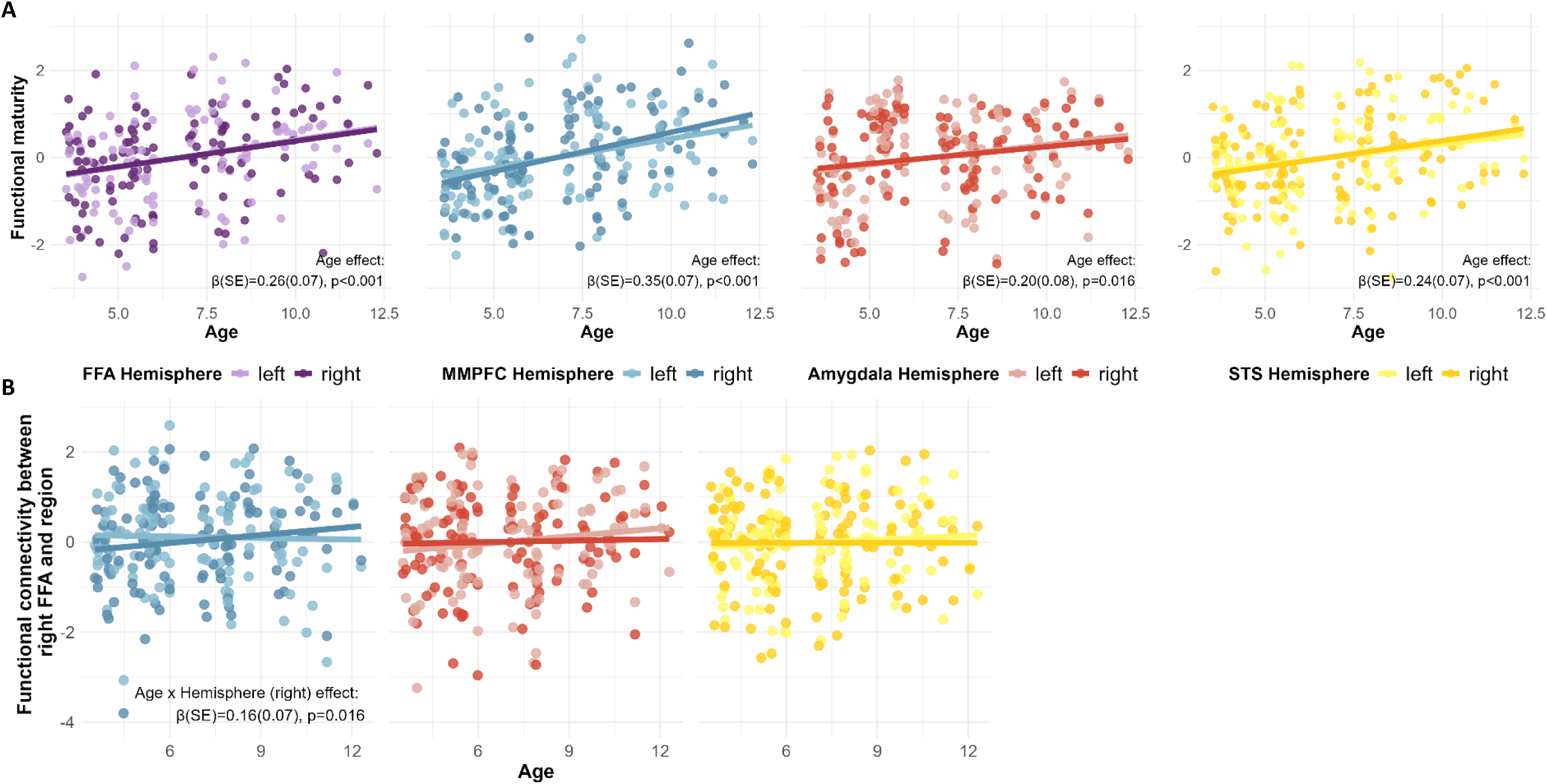
A) Association between age (x axis) and functional maturity (y axis) of FFA, MMPFC, amygdala, and STS. B) Association between age (x axis) and functional connectivity between right FFA and MMPFC, right FFA and amygdala, or right FFA and STS (y axis). Blue indicates relationships with MMPFC, red indicates relationships with amygdala, yellow indicates relationships with STS. Lighter colours correspond to the left hemisphere, darker colours correspond to the right hemisphere. In all scatterplots, lines represent linear regression fits estimated using the least-squares method; significant main effects are noted. N=117. FFA = Fusiform Face Area; MMPFC = Middle Medial Prefrontal Cortex, STS = Superior Temporal Sulcus; SE = Standard Error.

**Figure 3.**
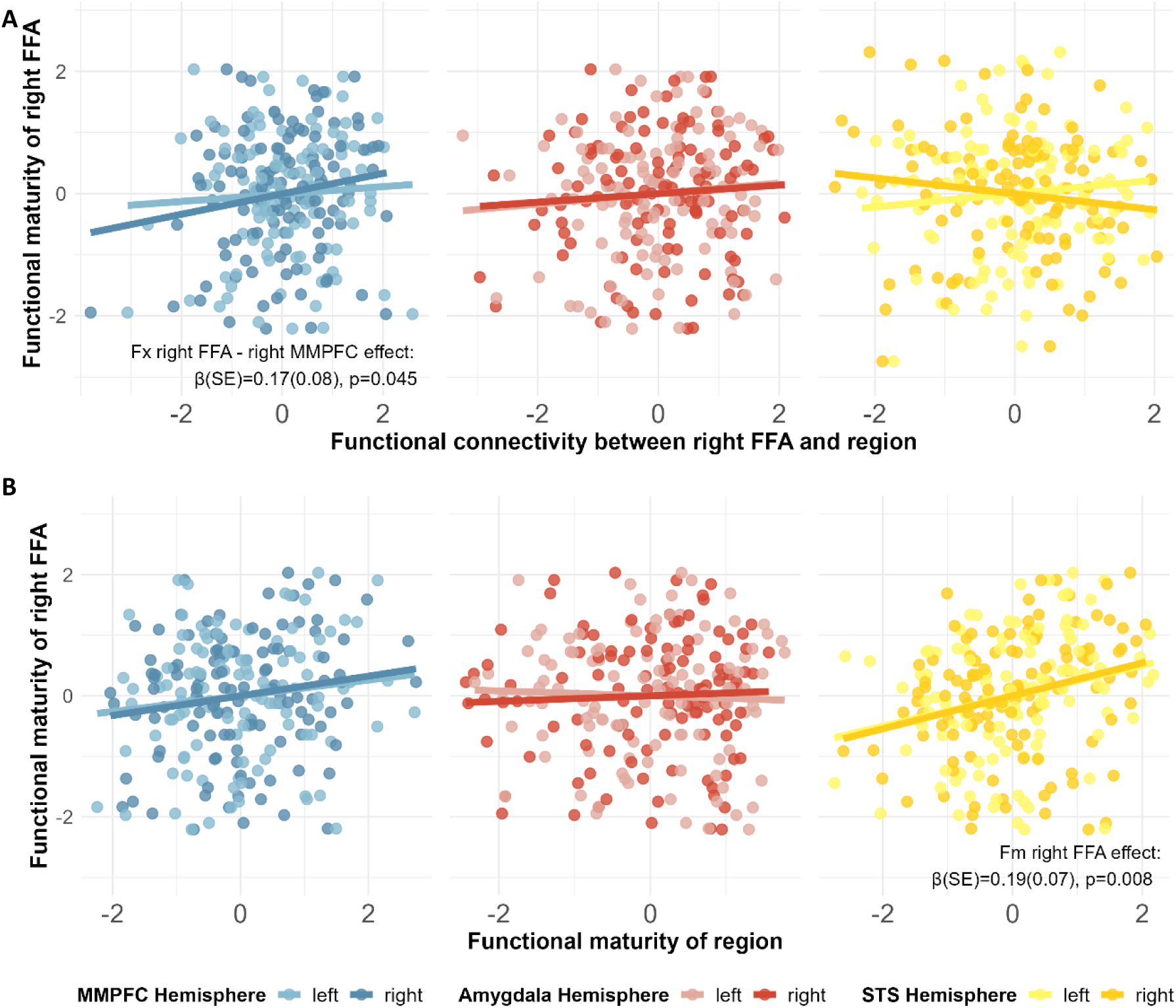
A) Association between functional connectivity (x axis) of right FFA and MMPFC (blue), amygdala (red), or STS (yellow) and functional maturity of right FFA (y axis). B) Association between functional maturity (x axis) of MMPFC (blue), amygdala (red), and STS (yellow) and functional maturity of right FFA (y axis). Blue indicates relationships with MMPFC, red indicates relationships with amygdala, yellow indicates relationships with STS. Lighter colours correspond to the left hemisphere, darker colours correspond to the right hemisphere of MMPFC, amygdala, and STS. In all scatterplots, lines represent linear regression fits estimated using the least-squares method; significant main effects are noted. N=117. FFA = Fusiform Face Area; MMPFC = Middle Medial Prefrontal Cortex; STS = Superior Temporal Sulcus; Fm = Functional Maturity; Fx = Functional Connectivity; SE = Standard Error.

Because the open dataset contains only one run of movie-viewing per participant, we used a split-half fROI definition approach (Kamps, Richardson et al., 2022): subject-specific fROIs were defined in one movie half (using one half of the reference timecourses) and response timecourses used for analyses were extracted in the opposite half; this procedure was iterated such that response timecourses from both movie halves were analysed.

Left and right amygdala ROIs were defined anatomically within individual participants using automatic segmentation by FreeSurfer Aseg Atlas (Fischl et al., 2002) within fMRIPrep.

#### Timecourse extraction

Preprocessed, smoothed timecourses were high-pass filtered (0.01Hz), outlier volumes were interpolated with cubic spline interpolation and censored, and the top five aCompCor were regressed out. Timecourses were then averaged across voxels within an ROI and z-scored, resulting in one timecourse per subject-specific ROI.

For fROIs (left and right FFA, MMPFC and STS), timecourses for each movie half were extracted from the fROI defined in the other, independent half of the movie; this approach ensures independence between data used to define an fROI and the timecourse data extracted and analysed. The middle 8s of the timecourse were excluded to prevent temporal autocorrelation across movie halves; note that the number of seconds excluded was increased from the preregistered value (6s; https://osf.io/hytsr) to obtain two movie halves of equal duration (given TR=2s).

For anatomically-defined left and right amygdala ROIs, full movie response timecourses were extracted and analysed.

#### Functional maturity

Functional maturity was defined as the z-scored Pearson correlation between each child’s timecourse and the average adult timecourse per ROI. For FFA, MMPFC and STS, the reference average adult timecourses were obtained from localiser-defined fROIs (FFA and STS: n=13 adults, available at https://osf.io/7a8w5/; MMPFC: n=17 adults, available at https://osf.io/mxkag/). Functional maturity in these regions was calculated separately for the first and second halves of the movie, and the two values were averaged to produce a single measure per ROI. For amygdala, the reference average adult timecourses were based on anatomically defined ROIs (n=33 adults), and functional maturity was computed using the entire movie.

#### Response magnitude to faces and scenes

Since functional maturity does not capture information about responses to faces per se, we also measured the magnitude of response to face events in our ROIs. Scene events were used as a negative control (i.e., dispreferred stimulus). We extracted z-scored response magnitude values from the peak timepoint of scenes that evoked reliable, positive responses from FFA (face events) or parahippocampal place area (PPA, scene events) in adults – as defined in prior research using the same adult sample studied here (n=33; Kamps, Richardson et al., 2022). There were 12 face events/timepoints and 12 scene events/timepoints. The peak timepoint was defined as the single timepoint (1 TR=2s) that evoked the highest average response in adults.

#### Inter-region correlations (i.e., functional connectivity)

For each child participant, inter-region correlations (IRCs) were defined as the z-scored Pearson correlations between each ROI timecourse. For IRCs involving fROIs (left and right FFA, left and right MMPFC, and left and right STS), IRCs were calculated per movie half and averaged to obtain one measure per ROI. For IRCs only involving amygdala ROIs, IRCs were calculated using the full movie response timecourse.

### 6.5 Statistical analyses

Data analyses were conducted and plots generated using R, version 4.4.3 (R Core Team, 2023).

#### Developmental change in functional responses of FFA, MMPFC, amygdala and STS

We tested for an effect of age on the functional maturity of each ROI using linear mixed-effects models with age, hemisphere, motion (mean FD) and an age-by-hemisphere interaction as fixed effects, alongside a random intercept for participant.

We also tested if the magnitude of response to face events in each ROI correlated with age using linear mixed-effects models. Peak magnitude for events per ROI were used as the outcome; one regression was conducted per event. Kamps, Richardson et al. (2022) identified 12 face events and 12 scene events (F01–F12 and S01–S12, respectively, following their numbering convention). We tested for fixed effects of age, hemisphere, motion and an age-by-hemisphere interaction effect, as well as a random effect of participant. We conducted a regression per event, rather than testing for effects on the average response magnitude across events per event type, because some events may be more sensitive to age-related differences than others and averaging across events could obscure age effects rather than increase power. We did not use mixed-effects models including responses to all events for similar reasons and because we did not expect to be sensitive to event-by-age interaction effects. Given the number of events and tests, results were corrected for multiple comparisons using a Bonferroni approach: effects were considered significant at α=0.004 for face and scene events (p=0.05/12, for 12 events per ROI).

We also investigated if functional connectivity between ROIs associated with age. To assess age effects on functional connectivity, we used a combination of linear models for within-region connectivity and linear mixed-effects models for between-region connectivity, with functional connectivity as the outcome in all models. For within-region connectivity (e.g., between left and right FFA ROIs), we tested for fixed effects of age and motion. For between-region connectivity (e.g., between right FFA and left/right MMPFC ROIs or right FFA and left/right amygdala ROIs), we tested for fixed effects of age, hemisphere, motion and an age-by-hemisphere interaction, alongside a random intercept for participant. Three primary models included connectivity of right FFA as the dependent variable; six secondary models included connectivity of left FFA, left and right MMPFC and amygdala as the dependent variable.

After conducting preregistered analyses, we conducted the same analyses described above with responses in STS. That is, we tested for age effects on functional maturity of STS, whether responses in STS to face and scene events associated with age, and whether functional connectivity between right FFA (primary models) and other ROIs (secondary models) and left and right STS increased with age.

#### Associations between functional maturity of FFA and its functional connectivity to MMPFC, amygdala and STS

A key hypothesis we were interested in testing in this dataset was whether functional maturity of FFA correlated with its functional connectivity to MMPFC or amygdala (https://osf.io/hytsr). Our preregistered primary analysis used linear mixed-effects models with functional maturity of right FFA as the dependent variable, functional connectivity between right FFA and the MMPFC, between right FFA and amygdala, participant’s age, hemisphere (left or right for MMPFC and amygdala), motion and an age-by-hemisphere-by-ROI interaction (and all embedded two-way interactions) as fixed effects, alongside a random effect of participant. As a secondary analysis, we preregistered a similar model with functional maturity of left FFA as the dependent variable. However, these models did not adequately converge and may have led to unstable estimates.

Instead, two separate, non-preregistered linear mixed-effects models were conducted to examine the association between right FFA functional maturity and each functional connectivity measure of interest. In each regression, functional connectivity between (1) right FFA and right or left MMPFC, (2) right FFA and right or left amygdala was the dependent variable with fixed effects of functional maturity of right FFA, age, motion and an age-by-hemisphere-by-functional maturity interaction (and all embedded two-way interactions), and a random intercept for participant. All non-significant interaction terms were removed from the final regression. Secondary models tested the same relationships for functional connectivity of left FFA to other ROIs and functional maturity of left FFA. In these models, the main dependent and independent variables were reversed relative to the preregistered plan because it is not possible to run a regression with repeated measures within a predictor (i.e., functional connectivity, which included a value per hemisphere, could not be an independent variable). Because all tests and claims are correlational, this necessary reversal does not impact our conclusions. Given the hypotheses we set out to test, we also conducted a non-preregistered analysis testing for a hemisphere-specific association – i.e., between right FFA functional maturity and its functional connectivity specifically to right MMPFC. We used a linear model with functional maturity of right FFA as dependent variable and fixed effects of functional connectivity between right FFA and right MMPFC, age, motion and an age-by-hemisphere-by-functional connectivity interaction (and all embedded two-way interactions).

We also ran non-preregistered linear mixed-effects models testing the same kind of associations, in the same way, using connectivity between right FFA and face-selective STS as dependent variable. The STS contains an early-developing face-selective region that could also plausibly shape or codevelop with FFA (Anzellotti et al., 2017, Deen et al., 2017, Kosakowski et al., 2024). Secondary models tested the same relationships for functional connectivity between left FFA and STS and functional maturity of left FFA instead of right FFA.

In additional non-preregistered analyses, we examined associations between the functional maturity of the right FFA, MMPFC, amygdala, and STS. We repeated the three linear mixed-effect models used previously, this time including the functional maturity of the MMPFC, amygdala, and STS as dependent variables. Secondary models tested the same relationships with functional maturity of left FFA as the independent variable.

#### Preregistered analyses of lateralisation of FFA face-response and its correlates

We also preregistered analyses to test associations between lateralisation of the FFA face-response and functional connectivity with or maturity of MMPFC and amygdala (https://osf.io/hytsr). Our lateralisation index was defined using the relative response to face and scene events, per hemisphere. However, given the use of a movie stimulus, these events cannot provide a clear estimate of face-selective responses - because almost all of the movie, including the scene events, involves faces being presented on the screen. Because of this, we were not confident in our lateralisation index and so include detailed methods and results in supplementary materials only (**Appendix S3**).

## 7. Results

### 7.1 Development of FFA face response and functional connectivity

Older children had more functionally mature (i.e., adult-like) FFA responses, controlling for motion (age effect: β(SE)=0.26(0.07), p=1.836×10^-4^, **Table S2**, **Figure 1**, Error! Reference source not found.**A**). Similar age effects on functional maturity were observed in MMPFC (β(SE)=0.35(0.07), p=4.841×10^-6^), amygdala (β(SE)=0.20(0.08), p=0.016, **Table S2**, **Figure 1**, Error! Reference source not found.**A**), and STS (β(SE)=0.24(0.07), p=3.968×10^-4^, **Table S2**, **Figure 1**, Error! Reference source not found.**A**).

FFA response magnitude did not vary with age for any face or scene event (defined in adults), controlling for motion and correcting for multiple comparisons (α=0.004 for face and scene events, tested separately; **Figure 1A**). However, MMPFC response magnitude decreased with age for one face event (F05; β(SE)=-0.23(0.06), p=3.798×10⁻⁴; see asterisk in **Figure 1B**, and **Figure S1** for interpretation), depicting Gus removing porcupine spikes from Peck’s head (evoking physical pain). In contrast, STS response magnitude increased with age for another face event (F10; β(SE)=0.14(0.04), p=7.311×10⁻⁴; see asterisk in **Figure 1D**, and **Figure S1**), showing Gus scolding the crocodile. No other regions showed significant age effects for face or scene events. Full results for all events and ROIs are provided in **Table S3**.

Functional connectivity between right FFA and the other regions of interest were generally positive across all children (right FFA and right MMPFC: M(SD) Pearson’s correlation value=0.19(0.23), t=8.73, p=2.193x10⁻^1^⁴; right FFA and left MMPFC: M(SD) Pearson’s correlation value=0.11(0.24), t=4.91, p=3.049x10^⁻6^; right FFA and right amygdala: M(SD) Pearson’s correlation value=0.34(0.19), t=19.05, p=1.654×10⁻^37^; right FFA and left amygdala: M(SD) Pearson’s correlation value=0.23(0.21), t=11.89, p=8.178×10⁻^22^, right FFA and right STS: M(SD) Pearson’s correlation value=0.38(0.20), t=20.41, p=3.353×10⁻^40^; right FFA and left STS: M(SD) Pearson’s correlation value=0.17(0.20), t=9.03, p=4.463×10⁻^15^; all t-tests against zero, non-preregistered, **Figure S2**). While functional connectivity between the right FFA and bilateral MMPFC did not increase with age (β(SE)=-0.01(0.09), p=0.873, **Table S4, Figure 2B, S2**), there was a significant age-by-hemisphere interaction such that connectivity between right FFA and right MMPFC increased more with age than connectivity between right FFA and left MMPFC (β(SE)=0.16(0.07), p=0.016, **Table S4, Figure 2B, S2**). In secondary analyses, connectivity between the right MMPFC and bilateral STS increased with age (β(SE)=0.15(0.07), p=0.038, **Table S4, Figure S2**). There were no other age effects/interactions on functional connectivity between left and right FFA, MMPFC, amygdala, and STS (all ps≥0.127, **Table S4, Figure 2B, S2, S3**).

### 7.2 Functional maturity of FFA: Associations with functional connectivity with or functional maturity of MMPFC, amygdala, or STS

We did not observe an association between more functionally mature (i.e., ‘adult-like’) right FFA responses and stronger functional connectivity between the right FFA and bilateral MMPFC or amygdala (all ps≥0.060, **Table 1A**, **Figure 3A**), or (in non-preregistered analyses) bilateral STS (p=0.106, **Table 1A**, **Figure 3A).** Because connectivity between right FFA and right MMPFC increased more with age than connectivity between right FFA and left MMPFC and including left MMPFC could obscure right-hemisphere-specific associations between the functional maturity of the right FFA and its connectivity to MMPFC, we conducted post-hoc analyses specifically testing for an association within the right hemisphere. In this analysis, children with more functionally mature right FFA responses had stronger functional connectivity between the right FFA and right MMPFC (β(SE)=0.17(0.08), p=0.045, **Table 1B**).

**Table 1.** Associations between functional maturity of right FFA and functional connectivity between FFA, MMPFC, amygdala, and STS.

| **A** |  |  |  |  |
| --- | --- | --- | --- | --- |
| **Fx right FFA - MMPFC** |  |  |  |  |
| Predictors | $\beta$ | SE | z | p |
| (Intercept) | 0.11 | 0.09 | 1.24 | 0.216 |
| Fm right FFA | 0.18 | 0.09 | 1.90 | 0.060 |
| MMPFC hemisphere (right) | -0.09 | 0.07 | -1.35 | 0.179 |
| Age | -0.06 | 0.09 | -0.62 | 0.536 |
| Mean FD | 0.20 | 0.09 | 2.15 | **0.034** |
| Age x Hemisphere (right) | 0.16 | 0.07 | 2.44 | **0.016** |
| Random effects | $\sigma$ | | | |
| ID | 0.84 |  |  |  |
| Residual | 0.51 |  |  |  |
| **Fx right FFA - amygdala** |  |  |  |  |
| Predictors | $\beta$ | SE | z | p |
| (Intercept) | -0.01 | 0.10 | -0.07 | 0.941 |
| Fm right FFA | 0.09 | 0.10 | 0.91 | 0.366 |
| Amygdala hemisphere (right) | 0.01 | 0.08 | 0.11 | 0.909 |
| Age | 0.06 | 0.09 | 0.70 | 0.487 |
| Mean FD | 0.06 | 0.09 | 0.63 | 0.532 |
| Random effects | $\sigma$ | | | |
| ID ($\sigma$) | 0.84 | | | |
| Residual ($\sigma$) | 0.63 | | | |
| **Fx right FFA - STS** |  |  |  |  |
| Predictors | $\beta$ | SE | z | p |
| (Intercept) | -0.01 | 0.09 | -0.14 | 0.890 |
| Fm right FFA | 0.12 | 0.08 | 1.63 | 0.106 |
| STS hemisphere (right) | -0.01 | 0.12 | -0.08 | 0.940 |
| Age | 0.02 | 0.07 | 0.23 | 0.817 |
| Mean FD | 0.19 | 0.07 | 2.65 | **0.009** |
| Random effects | $\sigma$ | | | |
| ID ($\sigma$) | 0.36 | | | |
| Residual ($\sigma$) | 0.90 | | | |
| **B** |  |  |  |  |
| **Fm right FFA** |  |  |  |  |
| Predictors | $\beta$ | SE | z | p |
| (Intercept) | 0.00 | 0.08 | -0.05 | 0.963 |
| Fx right FFA - right MMPFC | 0.17 | 0.08 | 2.03 | **0.045** |
| Age | 0.22 | 0.08 | 2.61 | **0.010** |
| Mean FD | -0.36 | 0.08 | -4.28 | **3.863 x 10<sup>-5</sup>** |

In secondary analyses of left FFA, children with more functionally mature left FFA responses had weaker functional connectivity between the left FFA and bilateral STS (β(SE)=-0.14(0.07), p=0.046, **Table S5**, **Figure S4A**).

In non-preregistered analyses, we did not observe evidence that children with more functionally mature right FFA responses had more functionally mature bilateral MMPFC or amygdala responses (all ps≥0.544, **Table 2**, **Figure 3B**), controlling for age. However, functional maturity of right FFA associated with functional maturity of bilateral STS, controlling for age (β(SE)=0.19(0.07), p=0.008, **Table 2**, **Figure 3B**). The same pattern was observed for the left FFA (**Table S6**, **Figure S4B**) – providing some evidence that our analyses were sensitive to associations between functional maturity of different regions.

**Table 2.**
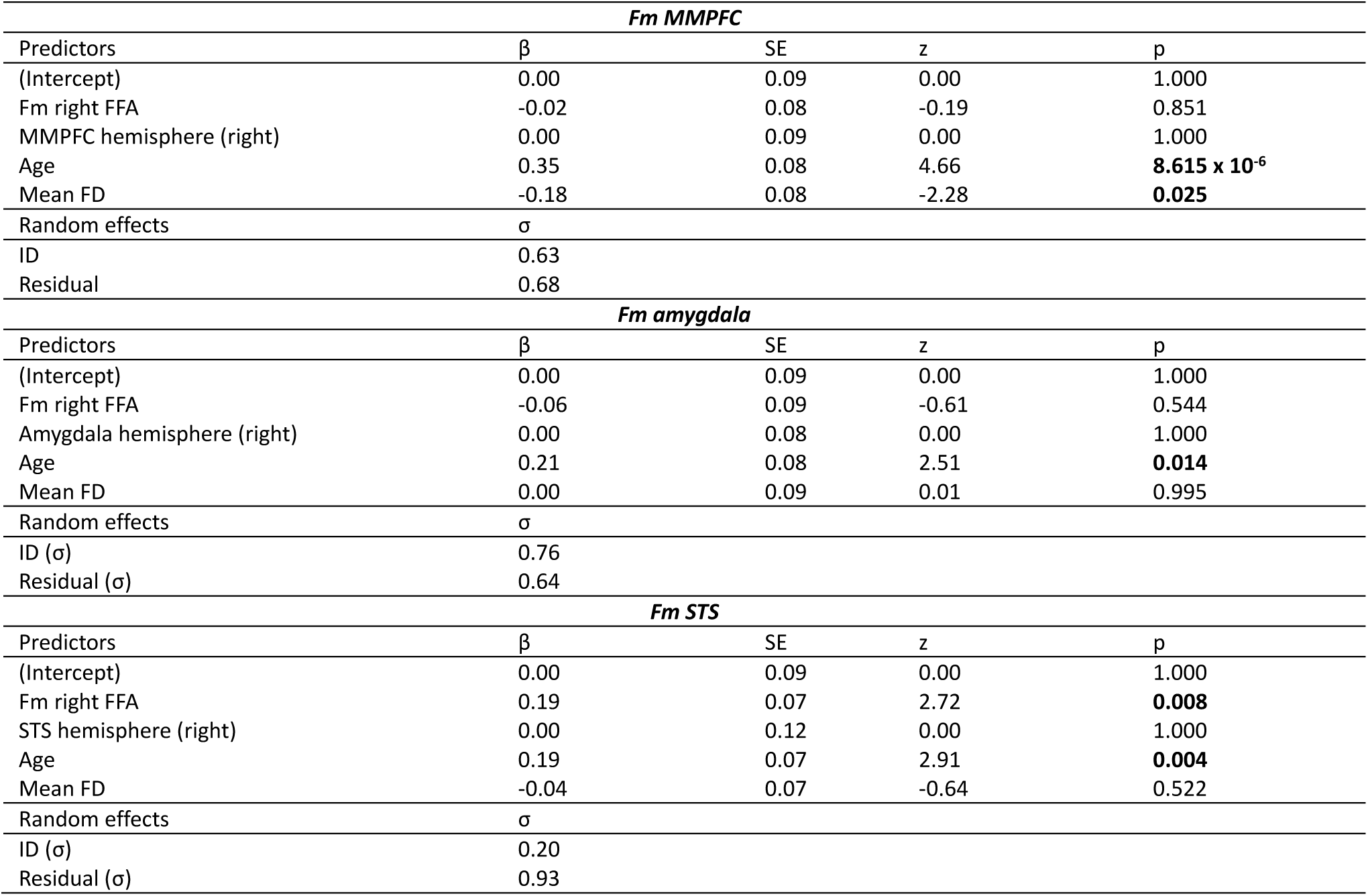
Associations between functional maturity of right FFA and functional maturity of MMPFC, amygdala, and STS.

Correlations among functional connectivity measures and among functional maturity measures are presented in **Figures S5** and **S6**, respectively.

## 8. Discussion

Evidence from fMRI studies with awake infants (Deen et al., 2017, Kosakowski et al., 2024) has fostered debate concerning drivers of face-selective responses in the right FFA (Livingstone et al., 2017, Op de Beeck et al., 2019, Powell et al., 2018). Here, we conducted opportunistic analyses on an existing cross-sectional paediatric fMRI dataset to test predictions of two non-mutually exclusive hypotheses articulated by Powell et al. (2018): that face-selective responses in right FFA develop alongside (1) higher-order brain systems involved in social interaction – and in particular, the MMPFC; and/or (2) subcortical regions which plausibly contain a face template – i.e., the amygdala. To test these hypotheses, we examined whether the functional connectivity between the FFA and the MMPFC or amygdala correlated with the functional maturity of right FFA. Children with stronger functional connectivity between the right FFA and right MMPFC had more functionally mature right FFA responses during movie-viewing. After conducting analyses focusing on MMPFC and amygdala, we extended our approach to study associations between FFA and STS; in doing so, we observed that children with more functionally mature STS responses had more functionally mature FFA responses, bilaterally. All analyses controlled for effects of age. Together, this evidence suggests that FFA co-develops with higher-order ‘social’ brain regions, including MMPFC and STS.

Our results are compatible with MMPFC (and STS) shaping the continued development of FFA responses in childhood – they do not speak directly to whether these regions play an early role in the initial development of face-selective responses in FFA. Opportunistic analyses of existing datasets are, by definition, not optimised for testing hypotheses other than those that the researchers had in mind when initially acquiring the data - and so should often be considered an initial step in a line of research. Our results and interpretations are limited in several ways, given the opportunistic nature of this study. For example, given that our analyses were conducted with a cross-sectional dataset of 3–12-year-old children, associations could reflect directional relationships, whereby MMPFC/STS play a causal role in the development of face responses in FFA, or vice-versa, or could simply reflect co-development of functional connectivity and maturity of these regions (beyond common effects of age). Further, associations between these different metrics could reflect a historical relationship: i.e., these measures would explain the initial development of face-selective FFA responses early in life, or they could simply reflect concurrent development among older children. More studies are needed to tease apart these alternatives, but such research will be challenging. For example, a micro-genetic study could measure these aspects of brain development concurrently as face-selective responses emerge (Dehaene-Lambertz et al., 2018) – but this would likely require multiple fMRI sessions per infant within the first months of life (Kosakowski et al., 2024, O’Doherty et al., 2026). Causal relationships between these different aspects of brain development could be informed through controlled rearing studies with non-human primate infants (Livingstone et al., 2017) or intervention studies investigating effects of positive, meaningful social interactions on attention to faces (Chawarska et al., 2025, Hayward et al., 2018) and FFA development. It may also be important for future research to employ interactive designs (Piazza et al., 2020) to measure brain responses related to self-relevant, socially meaningful interactions. The results of our opportunistic study help to constrain hypotheses and motivate experimental design and analytic choices for this future ‘planned’ research.

To test for an association between ‘higher order’ social brain regions and FFA development, our preregistered analyses focused on MMPFC, given its early involvement in processing self-relevant social interactions (Grossmann, 2025, Saxe and Kosakowski, 2025) and evidence for a face-selective response in MMPFC in infants (Deen et al., 2017, Kosakowski et al., 2024). We ultimately also conducted unplanned analyses with STS to build on prior relevant research in this area. In adults, both MMPFC and STS exert “top-down” influences on FFA responses (Anzellotti et al., 2017). The STS contains an early-developing face-selective subregion (Deen et al., 2017, Kosakowski et al., 2024), which (at least in adults) is particularly responsive to dynamic (i.e., moving) faces (Pitcher et al., 2011). The face-selective subregion of STS appears to be distinct from the subregion involved in social interaction perception (Deen et al., 2015, Masson and Isik, 2021, Im et al., 2025) and has been suggested to be important for facial expression recognition (Sliwinska and Pitcher, 2018). In children, prolonged development in face-selective STS (Kamps, Richardson et al., 2022, Tansey et al., 2023), entails increased functional correlations with higher-order brain regions involved in ‘theory of mind’ reasoning, including MMPFC (Kamps, Richardson et al., 2022). Our results suggest that this prolonged development also correlates with development in FFA. We also observed that children with more ‘adult-like’ *left* FFA responses had *weaker* functional connectivity between left FFA and STS; based on prior research, we suspect that this reflects the development of non-face-related responses in left FFA (i.e., visual word form area development) with age (Centanni et al., 2018, Behrmann et al., 2025).

Importantly, our positive evidence for an association between right FFA development and its functional connectivity with right MMPFC (and functional maturity of STS) does not diminish the potential (early or concurrent) role of the amygdala in FFA development. Null results are always difficult to interpret. Interestingly, functional connectivity was numerically stronger between right FFA and amygdala, relative to FFA and MMPFC (and strongest between right FFA and STS). It is possible that the associations we observed between FFA development and its connectivity to MMPFC and STS development reflect a prolonged developmental trajectory of MMPFC and STS and our sensitivity to developmental differences/variability in these measures, given the age range of our sample. Additional research is necessary to replicate and understand our pattern of evidence.

Additionally, our results do not provide evidence against a third hypothesis about the developmental origins of face-selective FFA responses, which is that they emerge through the repeated co-activation of neurons tuned to the low-level visual image statistics of faces (e.g., curvilinearity, foveal input and low spatial frequency (Livingstone et al., 2017, Arcaro et al., 2017, Powell et al., 2018). One challenge to this hypothesis raised by prior research is that this account, on its own, would predict bilateral face-selective FFA responses – and so it is difficult to resolve with known right-lateralised face responses in FFA (Powell et al., 2018). However, it remains unclear if early face-selective FFA responses are right-lateralised (Kosakowski et al., 2024), highlighting the possibility that neural mechanisms that drive the right-lateralisation of face responses – potentially including functional connectivity with higher-order brain regions – may be different from those that support the initial development of face-selective responses in FFA. Our experimental design did not enable us to confidently measure the lateralisation of face-selective responses (**Appendix S3**; though note that, consistent with Lesinger et al, 2023, functional connectivity between FFA and MMPFC, amygdala, and STS was numerically larger – i.e., stronger – between right hemisphere regions). Future research on the developmental trajectory of FFA face-response lateralisation, and the developmental association between lateralisation and selectivity, is needed. More broadly, it is likely that multiple factors and biases shape the development of face-selective FFA responses and all three hypotheses are worth interrogating in future research.

In conclusion, face-selective responses in FFA appear to be correlated with development of higher-order brain regions including MMPFC and STS in 3–12-year-old children. Future studies using longitudinal and intervention designs, socially relevant, self-referential stimuli, and experimental designs that allow for measuring the lateralisation of face-selective responses earlier in development are needed to clarify the mechanisms underlying the emergence and right-lateralisation of face-selective responses in the FFA.

## 9. Acknowledgements

The authors are grateful to the families who participated in this research.

## 12. Data and code availability statement

The movie fMRI data analysed during the current study (originally collected by Richardson et al., 2018) are publicly available on OpenNeuro (https://openneuro.org/datasets/ds000228). Further details about the fMRI analysis pipeline, and publicly available code, can be found at https://github.com/hrichardsonlab/fmri-analysis. Average timecourses from the referent adult populations, used as regressors in this study, are available through OSF (https://osf.io/7a8w5/ for FFA and LOC, and https://osf.io/mxkag/ for MMPFC and S2).

Scripts and data used for this analysis can be accessed at https://github.com/LorenaJS/FRIC.

## Abbreviations

EEG: Electroencephalography
FFA: Fusiform face area
FD: Framewise displacement
Fm: Functional maturity
fMRI: Functional magnetic resonance imaging
fNIRS: Functional near-infrared spectroscopy
Fx: Functional connectivity
M: Mean
(M)MPFC: (Middle) medial prefrontal cortex
LOC: Lateral occipital cortex
PPA: Parahippocampal place area
ROI: Region of interest
S2: Secondary somatosensory cortex
SD: Standard deviation
SE: Standard error
STS: Superior Temporal Sulcus
TR: Repetition time

