## Supplementary Material for "Fusiform face area development correlates with development in higher-order social brain regions"

**Appendix S1:** Pre-processing procedure by fMRIPrep 24.0.0

**Appendix S2:** Adult reference timecourses used for functional ROI definition

**Appendix S3:** Associations between lateralisation of FFA response and functional connectivity with/maturity of MMPFC, amygdala and STS

**Table S1:** Deviations from preregistered analysis plan

**Table S2:** Age effects on functional maturity of FFA, MMPFC, amygdala and STS

**Table S3:** Age effects on response magnitude of FFA, MMPFC, amygdala and STS to face and scene events

**Table S4:** Age effects on functional connectivity between FFA, MMPFC, amygdala and STS

**Table S5:** Associations between functional maturity of left FFA and its functional connectivity with MMPFC, amygdala and STS

**Table S6:** Associations between functional maturity of left FFA and functional maturity of MMPFC, amygdala and STS

**Table S7:** Age effects on FFA response lateralisation

**Table S8:** Associations between lateralisation of FFA response and functional connectivity between right FFA, MMPFC, amygdala and STS

**Table S9:** Associations between lateralisation of FFA response - calculated at $p<0.05$ - and functional connectivity between right FFA, MMPFC, amygdala and STS

**Table S10:** Associations between lateralisation of FFA response and functional connectivity between left FFA, MMPFC, amygdala and STS

**Table S11:** Associations between lateralisation of FFA response and functional maturity of FFA, MMPFC, amygdala, and STS

**Figure S1:** Significant associations between age and response magnitude of fROIs to face and scene events

**Figure S2:** Inter-region correlations (i.e., functional connectivity)

**Figure S3:** Age and functional connectivity between left FFA, MMPFC, amygdala and STS

**Figure S4:** Functional maturity of left FFA and functional maturity of/connectivity with MMPFC, amygdala and STS

**Figure S5:** Correlations of functional connectivity between FFA, MMPFC, amygdala and STS

**Figure S6:** Correlations of functional maturity between FFA, MMPFC, amygdala and STS

**Figure S7:** Age and FFA response lateralisation

**Figure S8:** Lateralisation of FFA response and functional connectivity of right FFA to/maturity of MMPFC, amygdala, and STS

**Figure S9:** Lateralisation of FFA response - calculated at $p<0.05$ - and functional connectivity of right FFA to MMPFC, amygdala and STS

**Figure S10:** Lateralisation of FFA response and functional connectivity of left FFA to MMPFC, amygdala and STS

**References**

***Appendix S1:*** *Pre-processing procedure by fMRIPrep 24.0.0*

The description below is a lightly edited version of the automatically-generated text by fMRIprep; which is released for use under the CC0 licence.

The T1w image was corrected for intensity non-uniformity (INU) with N4BiasFieldCorrection (Tustison et al., 2010), distributed with ANTs 2.5.1 (RRID:SCR_004757; Avants et al., 2008), and used as T1w-reference throughout the workflow. The T1w-reference was then skull-stripped with a Nipype implementation of the antsBrainExtraction.sh workflow (from ANTs), using OASIS30ANTs as target template. Brain tissue segmentation of cerebrospinal fluid (CSF), white-matter (WM) and grey-matter (GM) was performed on the brain-extracted T1w using fast (FSL, RRID:SCR_002823; Zhang et al., 2002). Brain surfaces were reconstructed using recon-all (FreeSurfer 7.3.2, RRID:SCR_001847; Dale et al., 1999), and the brain mask estimated previously was refined with a custom variation of the method to reconcile ANTs-derived and FreeSurfer-derived segmentations of the cortical grey-matter of Mindboggle (RRID:SCR_002438; Klein et al., 2017). Volume-based spatial normalisation to one standard space (MNI152NLin2009cAsym) was performed through nonlinear registration with antsRegistration (ANTs 2.5.1), using brain-extracted versions of both T1w reference and the T1w template. The following template was were selected for spatial normalisation and accessed with TemplateFlow 24.2.0 (Ciric et al., 2022): ICBM 152 Nonlinear Asymmetrical template version 2009c (Fonov et al., 2009; RRID:SCR_008796; TemplateFlow ID: MNI152NLin2009cAsym). For each of the BOLD runs found per subject (across all tasks and sessions), the following preprocessing was performed. First, a reference volume was generated, using a custom methodology of fMRIPrep, for use in head motion correction. Head-motion parameters with respect to the BOLD reference (transformation matrices, and six corresponding rotation and translation parameters) are estimated before any spatiotemporal filtering using mcflirt (FSL; Jenkinson et al., 2002). The BOLD reference was then co-registered to the T1w reference using bbregister (FreeSurfer) which implements boundary-based registration (Greve and Fischl, 2009). Co-registration was configured with six degrees of freedom. Several confounding time-series were calculated based on the preprocessed BOLD: framewise displacement (FD), DVARS and three region-wise global signals. FD was computed using two formulations following Power (absolute sum of relative motions; Power et al., 2014), and Jenkinson (relative root mean square displacement between affines; Jenkinson et al., 2002). FD and DVARS are calculated for each functional run, both using their implementations in Nipype (following the definitions by Power et al. (2014). The three global signals are extracted within the CSF, the WM, and the whole-brain masks. Additionally, a set of physiological regressors were extracted to allow for component-based noise correction (CompCor; Behzadi et al., 2007). Principal components are estimated after high-pass filtering the preprocessed BOLD time-series (using a discrete cosine filter with 128s cut-off) for the two CompCor variants: temporal (tCompCor) and anatomical (aCompCor). tCompCor components are then calculated from the top 2% variable voxels within the brain mask. For aCompCor, three probabilistic masks (CSF, WM and combined CSF+WM) are generated in anatomical space. The implementation differs from that of Behzadi et al. in that instead of eroding the masks by 2 pixels on BOLD space, a mask of pixels that likely contain a volume fraction of GM is subtracted from the aCompCor masks. This mask is obtained by dilating a GM mask extracted from the FreeSurfer's aseg segmentation, and it ensures components are not extracted from voxels containing a minimal fraction of GM. Finally, these masks are resampled into BOLD space and binarised by thresholding at 0.99 (as in the original implementation). Components are also calculated separately within the WM and CSF masks. For each CompCor decomposition, the k components with the largest singular values are retained, such that the retained components' time series are sufficient to explain 50 percent of variance across the nuisance mask (CSF, WM, combined, or temporal). The remaining components are dropped from consideration. The head-motion estimates calculated in the correction step were also placed within the corresponding confounds file. The confound time series derived from head motion estimates and global signals were expanded with the inclusion of temporal derivatives and quadratic terms for each (Satterthwaite et al., 2013). Frames that exceeded a threshold of 0.5 mm FD or 1.5 standardised DVARS were annotated as motion outliers. Additional nuisance timeseries are calculated by means of principal components analysis of the signal found within a thin band (crown) of voxels around the edge of the brain, as proposed by Patriat et al. (2017). All resamplings can be performed with a single interpolation step by composing all the pertinent transformations (i.e., head-motion transform matrices, susceptibility distortion correction when available, and co-registrations to anatomical and output spaces). Gridded (volumetric) resamplings were performed using nitransforms, configured with cubic B-spline interpolation.

Many internal operations of fMRIPrep use Nilearn 0.10.4 (RRID:SCR_001362; Abraham et al., 2014), mostly within the functional processing workflow. For more details of the pipeline, see the section corresponding to workflows in fMRIPrep's documentation.

***Appendix S2:*** *Adult reference timecourses used for functional ROI definition*

To define functional ROIs, we used adult reference timecourses as regressors in first-level models of children's neural activity during the movie. Movie timecourses from localiser-defined FFA and LOC were originally extracted by Kamps, Richardson et al. (2022) and are publicly available (n=13 adults; independent from those in the open dataset; https://osf.io/7a8w5/). FFA, LOC, and STS ROIs were all defined within large search spaces previously defined in an independent sample of adults (Julian et al., 2012). Localiser-defined FFA and STS ROIs were the 100 most responsive voxels to a faces > objects contrast (Kanwisher et al., 1997). Lateral occipital cortex (LOC) ROIs were the 100 most responsive voxels to an objects > scrambled objects localiser contrast (Grill-Spector et al., 1998).

Movie timecourses from localiser-defined MMPFC and S2 were originally extracted by Richardson and colleagues in unpublished research and are shared on OSF (n=17 adults; all of whom are included in the open dataset; https://osf.io/mxkag/). Localiser-defined MMPFC ROIs were the 80 voxels within a large MMPFC search space (Dufour et al., 2013) most responsive to a false belief > false photograph localiser contrast (Dodell-Feder et al., 2011; https://saxelab.mit.edu/use-our-efficient-false-belief-localizer/). Secondary sensory motor cortex (S2) ROIs were the 80 voxels within left and right S2 search spaces (Bruneau et al., 2015) most responsive to a physical pain > emotional pain localiser contrast (https://saxelab.mit.edu/theory-mind-and-pain-matrix-localizer-narratives/).

***Appendix S3:*** *Associations between lateralisation of FFA response and functional connectivity with/maturity of MMPFC, amygdala and STS*

***S3.1*** *Methods*

***S3.1.1*** *Lateralisation index*

We calculated a lateralisation index (LI) for FFA responses by counting the number of voxels in each hemisphere within a large predefined group FFA ROI (Julian et al., 2012) that showed significant activation to the contrast of faces > scenes events in the movie. To ensure equivalent search volumes, the left FFA search space was defined by flipping the right FFA search space (available at https://osf.io/mxkag/). The LI was calculated as (Desmond et al., 1995):

LI = (Number of voxels in left FFA - Number of voxels in right FFA) / (Number of voxels in left FFA + Number of voxels in right FFA)

We preregistered calculating LI at two thresholds to ensure that effects were not threshold dependent: $p<0.001$ ($T>3.10$) and $p<0.01$ ($T>2.33$). However, at these significance thresholds only 9 (at $p<0.001$) and 44 (at $p<0.01$) participants had at least 20 suprathreshold voxels across left and right FFA - severely limiting the sample size for subsequent analyses. We accidentally omitted the 20 suprathreshold voxel criterion from our preregistration, but this criterion is common in these kinds of analyses to reduce impacts of noise inherent to a neural measure based on <20 voxels (Zimmermann et al., 2024, Kewenig et al., 2024, Bailey et al., 2025). Given the restricted sample size at the preregistered thresholds, we decided to conduct the preregistered LI analyses using more lenient (non-preregistered) thresholds: $p<0.05$ ($T>1.645$, yielding 97 children for analyses; M(SD)=6.75(2.32) years of age; 52 females) and $p<0.10$ ($T>1.289$, yielding 110 children for analyses; M(SD)=6.70(2.33) years of age; 58 females). We report results using a threshold of $p<0.10$ and note any differences observed when applying the more conservative $p<0.05$ threshold.

***S3.1.2*** *Statistical analyses*

***S3.1.2.1*** *Development of lateralisation of face responses in FFA*

We tested for an effect of age on face response lateralisation using linear models with motion as a fixed effect (non-preregistered).

***S3.1.2.2*** *Associations between lateralisation of FFA response and functional connectivity with/functional maturity of MMPFC, amygdala, and STS*

We tested whether functional connectivity between the FFA and the MMPFC, amygdala, and/or STS correlated with the response lateralisation. Preregistered analyses used linear mixed-effects models with LI as the dependent variable and functional connectivity between right FFA and MMPFC, right FFA and amygdala, age, hemisphere, and an age-by-hemisphere-by-ROI interaction (and all embedded two-way interactions) as fixed effects, as well as a random intercept for participant (with analogous secondary models for the left FFA). However, these models did not converge and could have led to unstable estimates. Instead, we ran and report two separate non-preregistered linear mixed-effects models with functional connectivity as the dependent variable and LI, age, hemisphere, motion and an age-by-hemisphere-by-LI interaction (and all lower-order terms) as fixed effects, with a random intercept for participant. Models were run for connectivity between (1) right FFA and right and left MMPFC and (2) right FFA and right and left amygdala; interaction terms were removed if non-significant. Secondary models tested the same relationships for the left FFA. Post-hoc, non-preregistered analyses used the same model structure to test for correlations between response lateralisation and functional connectivity between FFA and STS.

Second, we tested if functional maturity of the MMPFC and amygdala correlated with FFA response lateralisation. Preregistered models paralleled those above, substituting functional maturity for connectivity, but again did not converge. We therefore fit two mixed-effects models with functional maturity of the (1) MMPFC and (2) amygdala as the dependent variable and LI, age, hemisphere, motion and a hemisphere-by-LI interaction (and all embedded two-way interactions), alongside a random intercept for participant. Non-significant interactions were removed from the final regression. An additional non-preregistered model tested for an analogous correlation with functional maturity of the STS.

### *S3.2 Results*

#### *S3.2.1 Developmental change in LI*

FFA face responses were generally not lateralised across all children (LI M(SD)=0.06(0.51), t=1.20, p=0.232; t-test against zero, non-preregistered, N=110, age M(SD)=6.70(2.33)) and did not change with age (β(SE)=-0.16(0.10), p=0.096, **Table S7**, **Figure S7**); analyses at the p<0.05 threshold revealed the same pattern of results.

#### *S3.2.1 Associations between lateralisation of FFA response and functional connectivity with/functional maturity of MMPFC, amygdala, and STS*

Children with more functionally connected right FFA and bilateral MMPFC had a more left lateralised face response (β(SE)=0.22(0.10), p=0.027, **Table S8**, **Figure S8A**). There was also a lateralisation-by-hemisphere effect (β(SE)=-0.23(0.07), p=8.360 x $10^{-4}$, **Table S8**, **Figure S8A**), indicating that association was stronger for the left hemisphere of MMPFC; that is: children with higher functional connectivity between right FFA and left MMPFC had more left-lateralised face responses. While the data look similar at both thresholds for calculating LI (**Figures S8, S9**), these results were only statistically significant at the more lenient threshold (i.e., at p<0.10, N=110, but not p<0.05, N=97, **Tables S8, S9**).

The lateralisation of face responses in FFA did not associate with the functional connectivity between right FFA and amygdala or STS (all ps≥0.358, **Table S8**, **Figure S8A**). There were no significant main effects or interactions in secondary models with left FFA (**Table S10**, **Figure S10**). These primary and secondary analyses showed a similar pattern of results at the more conservative threshold (p<0.05).

There were no associations between functional maturity of MMPFC, amygdala, or STS and the lateralisation of face responses in FFA, controlling for age (all ps>0.185, **Table S11A**, **Figure S8B**). Further, functional maturity of FFA was not associated with the lateralisation of face responses in FFA, controlling for age (p=0.734, **Table S11B**). This pattern was observed in analyses using both thresholds for calculating LI.

### *S3.3 Discussion*

On the whole, we recommend caution in interpreting the results of the lateralisation analyses. Prior research characterising right-lateralisation primarily uses controlled experimental stimuli that enable comparing response magnitude across conditions (face and non-face), and comparing the size of the face-selective response across hemispheres (Kanwisher et al., 1997, Kosakowski et al., 2024, Liu et al., 2024, Thome et al., 2022). We preregistered a similar approach using ‘face’ and ‘scene’ events, which were defined as events that reliably drove responses in adult FFA and PPA, respectively. On reflection, we do not believe that contrasting these events provides a clean estimate of face-selective responses - because almost all of the movie, including the scene events, involves faces being presented on the screen. It is therefore hard to know if a (more) ‘left-lateralised’ face response reflects more face-selective voxels in response to face events in the left hemisphere, more face-selective voxels in response to scene events (which also have faces) in the right hemisphere, or both. The issues with our face > scene contrast may have contributed to the low number of participants with sufficient suprathreshold FFA voxels to take forward into analyses at our planned thresholds, which were fairly standard ($p<0.01$ and $p<0.001$). We decided to present these results because we preregistered the analysis and the question of response lateralisation – and its developmental trajectory – is of interest to the field and relevant to the hypotheses tested in this paper. We hope that by doing so, future researchers can consider these issues carefully in their own research.

***Table S1:*** *Deviations from preregistered analysis plan*

| ***Deviations from preregistered analysis plan*** | | |
|---|---|---|
| Section | Preregistered | Modification |
| fMRI analysis: motion treatment | Apply threshold of standardised DVARS>1.5 | Threshold not applied |
| fMRI analysis: timecourse extraction | Exclude middle 6 s of timecourses | Middle 8 s of timecourses excluded |
| Developmental change in functional responses of ROIs | ROIs: FFA, MMPFC and amygdala | ROIs: FFA, MMPFC, amygdala and STS |
| Associations between functional maturity of FFA and its functional connectivity to ROIs | ROIs: FFA, MMPFC and amygdala<br>Statistics: Full linear mixed-effects model including all connectivity measures | ROIs: FFA, MMPFC, amygdala and STS<br>Statistics: linear mixed-effects model reversing independent and dependent variable, one for each pair of ROIs |
| Associations between lateralisation of FFA face-response and functional connectivity with or maturity of ROIs | ROIs: FFA, MMPFC and amygdala<br>Lateralisation index of FFA: calculated at p<0.001, uncorrected (T>3.10) and p<0.01, uncorrected (T>2.33).<br>Statistics: Full linear mixed-effects model including all connectivity measures<br>Statistics: Full linear mixed-effects model including all maturity measures<br>Planned as one of the main dependent variables measuring FFA development | ROIs: FFA, MMPFC, amygdala and STS<br>Lateralisation index of FFA: calculated at p<0.05, uncorrected (T>1.645) and p<0.10, uncorrected (T>1.289).<br>Statistics: linear mixed-effects model reversing independent and dependent variable, one for each pair of ROIs<br>Statistics: linear mixed-effects model reversing independent and dependent variable, one for each pair of ROIs<br>Presented in supplementary materials only given perceived issues with contrast/experimental design |
| ***Additions to preregistered analysis plan*** | | |
| Section | Tests | |
| Developmental change in functional responses of other ROIs | Associations between age and lateralisation of FFA face-response<br>T-tests of functional connectivity or lateralisation index values against zero in the full sample | |
| Associations between functional maturity of FFA and its functional connectivity to other ROIs | Associations between functional maturity of right FFA and functional connectivity of right FFA and right MMPFC (linear model)<br>Associations between the functional maturity of the FFA, MMPFC, amygdala and STS (linear mixed-effects model reversing independent and dependent variable, one for each pair of ROIs) | |
| Associations between lateralisation of FFA face-response and functional connectivity with or maturity of other ROIs | Associations between lateralisation index (calculated at p<0.05 and p<0.10) and functional maturity of FFA (linear mixed-effects model reversing independent and dependent variable, one for each lateralisation index threshold) | |

*FFA = Fusiform Face Area; MMPFC = Middle Medial Prefrontal Cortex; STS = Superior Temporal Sulcus; ROI = Region of Interest.*

***Table S2:*** *Age effects on functional maturity of FFA, MMPFC, amygdala and STS*

| | | | | |
|---|---|---|---|---|
| | | ***FFA*** | | |
| Predictors | B | SE | z | p |
| (Intercept) | 0.00 | 0.09 | 0.00 | 1.000 |
| Age | 0.26 | 0.07 | 3.87 | **1.836 x $10^{-4}$** |
| Hemisphere (right) | 0.00 | 0.11 | 0.00 | 1.000 |
| Mean FD | -0.22 | 0.07 | -3.19 | **0.002** |
| Random effects | σ | | | |
| ID | 0.44 | | | |
| Residual | 0.83 | | | |
| | | ***MMPFC*** | | |
| Predictors | β | SE | z | p |
| (Intercept) | 0.00 | 0.09 | 0.00 | 1.000 |
| Age | 0.35 | 0.07 | 4.80 | **4.841 x $10^{-6}$** |
| Hemisphere (right) | 0.00 | 0.09 | 0.00 | 1.000 |
| Mean FD | -0.17 | 0.07 | -2.38 | **0.019** |
| Random effects | σ | | | |
| ID | 0.62 | | | |
| Residual | 0.68 | | | |
| | | ***Amygdala*** | | |
| Predictors | β | SE | z | p |
| (Intercept) | 0.00 | 0.09 | 0.00 | 1.000 |
| Age | 0.20 | 0.08 | 2.44 | **0.016** |
| Hemisphere (right) | 0.00 | 0.08 | 0.00 | 1.000 |
| Mean FD | 0.02 | 0.08 | 0.24 | 0.812 |
| Random effects | σ | | | |
| ID (σ) | 0.75 | | | |
| Residual (σ) | 0.64 | | | |
| | | ***STS*** | | |
| Predictors | β | SE | z | p |
| (Intercept) | 0.00 | 0.09 | 0.00 | 1.000 |
| Age | 0.24 | 0.07 | 3.66 | **3.968 x $10^{-4}$** |
| Hemisphere (right) | 0.00 | 0.12 | 0.00 | 1.000 |
| Mean FD | -0.11 | 0.07 | -1.63 | 0.098 |
| Random effects | σ | | | |
| ID (σ) | 0.26 | | | |
| Residual (σ) | 0.93 | | | |

*Age-by-hemisphere interaction terms were not statistically significant and were excluded from the final models. Reported βs are standardised, ps<0.05 are shown in bold and scientific notation is used for ps<0.001. N=117. FFA = Fusiform Face Area; MMPFC = Middle Medial Prefrontal Cortex; STS = Superior Temporal Sulcus; FD = Framewise Displacement; SE = Standard Error.*

***Table S3:*** *Age effects on response magnitude of FFA, MMPFC, amygdala and STS to face and scene events*

| | | ***FFA*** | | | |
|---|---|---|---|---|---|
| F01 | Predictors | β | SE | z | p |
| | (Intercept) | -0.21 | 0.08 | -2.66 | 0.008 |
| | Age | -0.09 | 0.06 | -1.36 | 0.176 |
| | Hemisphere (right) | -0.19 | 0.08 | -2.19 | 0.031 |
| | Mean FD | 0.00 | 0.07 | 0.07 | 0.944 |
| | Random effects | σ | | | |
| | ID | 0.49 | | | |
| | Residual | 0.61 | | | |
| F09 | Predictors | β | SE | z | p |
| | (Intercept) | -0.28 | 0.07 | -4.27 | **$3.074 \times 10^{-5}$** |
| | Age | -0.05 | 0.06 | -0.93 | 0.353 |
| | Hemisphere (right) | -0.25 | 0.08 | -3.20 | **0.002** |
| | Mean FD | 0.01 | 0.06 | 0.14 | 0.885 |
| | Random effects | σ | | | |
| | ID | 0.40 | | | |
| | Residual | 0.58 | | | |
| F11 | Predictors | β | SE | z | p |
| | (Intercept) | -0.40 | 0.06 | -6.38 | **$1.489 \times 10^{-9}$** |
| | Age | -0.04 | 0.05 | -0.68 | 0.500 |
| | Hemisphere (right) | 0.01 | 0.07 | 0.19 | 0.847 |
| | Mean FD | 0.09 | 0.05 | 1.72 | 0.088 |
| | Random effects | σ | | | |
| | ID | 0.42 | | | |
| | Residual | 0.50 | | | |
| F04 | Predictors | β | SE | z | p |
| | (Intercept) | -0.28 | 0.06 | -4.81 | **$3.183 \times 10^{-6}$** |
| | Age | -0.02 | 0.05 | -0.49 | 0.624 |
| | Hemisphere (right) | 0.04 | 0.06 | 0.63 | 0.531 |
| | Mean FD | 0.01 | 0.05 | 0.14 | 0.892 |
| | Random effects | σ | | | |
| | ID | 0.40 | | | |
| | Residual | 0.46 | | | |
| F10 | Predictors | β | SE | z | p |
| | (Intercept) | 0.41 | 0.05 | 7.72 | **$4.692 \times 10^{-13}$** |
| | Age | 0.09 | 0.04 | 2.23 | 0.028 |
| | Hemisphere (right) | -0.11 | 0.06 | -1.65 | 0.103 |
| | Mean FD | -0.08 | 0.04 | -1.85 | 0.067 |
| | Random effects | σ | | | |
| | ID | 0.30 | | | |
| | Residual | 0.49 | | | |
| F12 | Predictors | β | SE | z | p |
| | (Intercept) | 0.39 | 0.06 | 6.80 | **$9.641 \times 10^{-11}$** |
| | Age | 0.05 | 0.04 | 1.09 | 0.276 |
| | Hemisphere (right) | -0.04 | 0.07 | -0.56 | 0.578 |
| | Mean FD | -0.08 | 0.04 | -1.84 | 0.069 |
| | Random effects | σ | | | |
| | ID | 0.23 | | | |
| | Residual | 0.56 | | | |
| F07 | Predictors | β | SE | z | p |
| | (Intercept) | 0.05 | 0.07 | 0.72 | 0.471 |
| | Age | -0.01 | 0.06 | -0.19 | 0.853 |
| | Hemisphere (right) | -0.11 | 0.08 | -1.33 | 0.187 |
| | Mean FD | -0.09 | 0.06 | -1.50 | 0.136 |
| | Random effects | σ | | | |
| | ID | 0.41 | | | |
| | Residual | 0.59 | | | |
| F08 | Predictors | β | SE | z | p |
| | (Intercept) | -0.08 | 0.06 | -1.17 | 0.245 |
| | Age | -0.06 | 0.05 | -1.27 | 0.208 |
| | Hemisphere (right) | 0.04 | 0.08 | 0.51 | 0.611 |
| | Mean FD | -0.07 | 0.05 | -1.39 | 0.166 |

| | | | | | |
|---|---|---|---|---|---|
| | Random effects | σ | | | |
| | ID | 0.33 | | | |
| | Residual | 0.60 | | | |
| F06 | Predictors | β | SE | z | p |
| | (Intercept) | -0.04 | 0.06 | -0.59 | 0.555 |
| | Age | 0.05 | 0.05 | 1.03 | 0.306 |
| | Hemisphere (right) | -0.02 | 0.08 | -0.26 | 0.794 |
| | Mean FD | 0.07 | 0.05 | 1.27 | 0.208 |
| | Random effects | σ | | | |
| | ID | 0.31 | | | |
| | Residual | 0.60 | | | |
| F05 | Predictors | β | SE | z | p |
| | (Intercept) | -0.04 | 0.06 | -0.66 | 0.508 |
| | Age | 0.06 | 0.05 | 1.27 | 0.207 |
| | Hemisphere (right) | 0.08 | 0.08 | 0.99 | 0.325 |
| | Mean FD | -0.01 | 0.05 | -0.17 | 0.865 |
| | Random effects | σ | | | |
| | ID | 0.24 | | | |
| | Residual | 0.60 | | | |
| F02 | Predictors | β | SE | z | p |
| | (Intercept) | 0.46 | 0.06 | 7.32 | **$8.232 \times 10^{-12}$** |
| | Age | 0.05 | 0.05 | 0.95 | 0.344 |
| | Hemisphere (right) | -0.18 | 0.07 | -2.68 | 0.009 |
| | Mean FD | 0.00 | 0.06 | -0.07 | 0.947 |
| | Random effects | σ | | | |
| | ID | 0.44 | | | |
| | Residual | 0.49 | | | |
| F03 | Predictors | β | SE | z | p |
| | (Intercept) | 0.25 | 0.07 | 3.52 | **$5.417 \times 10^{-4}$** |
| | Age | 0.04 | 0.06 | 0.65 | 0.518 |
| | Hemisphere (right) | -0.11 | 0.08 | -1.45 | 0.151 |
| | Mean FD | 0.02 | 0.06 | 0.40 | 0.690 |
| | Random effects | σ | | | |
| | ID | 0.48 | | | |
| | Residual | 0.59 | | | |
| S07 | Predictors | β | SE | z | p |
| | (Intercept) | -0.21 | 0.08 | -2.66 | 0.008 |
| | Age | -0.09 | 0.06 | -1.36 | 0.176 |
| | Hemisphere (right) | -0.19 | 0.08 | -2.19 | 0.031 |
| | Mean FD | 0.00 | 0.07 | 0.07 | 0.944 |
| | Random effects | σ | | | |
| | ID | 0.49 | | | |
| | Residual | 0.61 | | | |
| S02 | Predictors | β | SE | z | p |
| | (Intercept) | -0.01 | 0.06 | -0.10 | 0.918 |
| | Age | 0.06 | 0.05 | 1.19 | 0.235 |
| | Hemisphere (right) | -0.23 | 0.07 | -3.56 | **$5.539 \times 10^{-4}$** |
| | Mean FD | 0.03 | 0.05 | 0.58 | 0.562 |
| | Random effects | σ | | | |
| | ID | 0.38 | | | |
| | Residual | 0.48 | | | |
| S03 | Predictors | β | SE | z | p |
| | (Intercept) | 0.22 | 0.05 | 4.10 | **$5.956 \times 10^{-5}$** |
| | Age | 0.02 | 0.04 | 0.44 | 0.659 |
| | Hemisphere (right) | -0.03 | 0.06 | -0.51 | 0.611 |
| | Mean FD | -0.05 | 0.04 | -1.03 | 0.305 |
| | Random effects | σ | | | |
| | ID | 0.31 | | | |
| | Residual | 0.48 | | | |
| S04 | Predictors | β | SE | z | p |
| | (Intercept) | 0.54 | 0.06 | 8.72 | **$2.184 \times 10^{-15}$** |
| | Age | -0.06 | 0.05 | -1.19 | 0.238 |
| | Hemisphere (right) | -0.08 | 0.07 | -1.14 | 0.256 |
| | Mean FD | 0.02 | 0.06 | 0.33 | 0.739 |

| | | | | | |
|---|---|---|---|---|---|
| | Random effects | σ | | | |
| | ID | 0.39 | | | |
| | Residual | 0.50 | | | |
| S12 | Predictors | β | SE | z | p |
| | (Intercept) | 0.44 | 0.06 | 7.34 | **8.288 x 10$^{-12}$** |
| | Age | 0.02 | 0.05 | 0.36 | 0.718 |
| | Hemisphere (right) | 0.00 | 0.06 | 0.00 | 0.998 |
| | Mean FD | -0.03 | 0.05 | -0.64 | 0.525 |
| | Random effects | σ | | | |
| | ID | 0.44 | | | |
| | Residual | 0.44 | | | |
| S11 | Predictors | β | SE | z | p |
| | (Intercept) | 0.18 | 0.06 | 3.00 | **0.003** |
| | Age | 0.02 | 0.05 | 0.40 | 0.689 |
| | Hemisphere (right) | 0.18 | 0.07 | 2.51 | 0.014 |
| | Mean FD | 0.00 | 0.05 | -0.07 | 0.946 |
| | Random effects | σ | | | |
| | ID | 0.36 | | | |
| | Residual | 0.53 | | | |
| S10 | Predictors | β | SE | z | p |
| | (Intercept) | -0.30 | 0.07 | -4.48 | **1.208 x 10$^{-5}$** |
| | Age | -0.05 | 0.05 | -1.04 | 0.303 |
| | Hemisphere (right) | 0.10 | 0.08 | 1.22 | 0.226 |
| | Mean FD | -0.01 | 0.05 | -0.25 | 0.803 |
| | Random effects | σ | | | |
| | ID | 0.33 | | | |
| | Residual | 0.61 | | | |
| S06 | Predictors | β | SE | z | p |
| | (Intercept) | -0.04 | 0.06 | -0.66 | 0.508 |
| | Age | 0.06 | 0.05 | 1.27 | 0.207 |
| | Hemisphere (right) | 0.08 | 0.08 | 0.99 | 0.325 |
| | Mean FD | -0.01 | 0.05 | -0.17 | 0.865 |
| | Random effects | σ | | | |
| | ID | 0.24 | | | |
| | Residual | 0.60 | | | |
| S08 | Predictors | β | SE | z | p |
| | (Intercept) | 0.23 | 0.06 | 3.87 | **1.487 x 10$^{-4}$** |
| | Age | -0.08 | 0.05 | -1.76 | 0.081 |
| | Hemisphere (right) | -0.11 | 0.07 | -1.68 | 0.096 |
| | Mean FD | -0.06 | 0.05 | -1.10 | 0.275 |
| | Random effects | σ | | | |
| | ID | 0.37 | | | |
| | Residual | 0.48 | | | |
| S05 | Predictors | β | SE | z | p |
| | (Intercept) | -0.13 | 0.06 | -2.09 | 0.038 |
| | Age | 0.00 | 0.04 | 0.11 | 0.911 |
| | Hemisphere (right) | -0.07 | 0.09 | -0.78 | 0.439 |
| | Mean FD | 0.05 | 0.05 | 0.98 | 0.326 |
| | Random effects | σ | | | |
| | ID | 0.00 | | | |
| | Residual | 0.66 | | | |
| S09 | Predictors | β | SE | z | p |
| | (Intercept) | 0.09 | 0.06 | 1.49 | 0.137 |
| | Age | 0.00 | 0.04 | 0.08 | 0.935 |
| | Hemisphere (right) | 0.03 | 0.08 | 0.42 | 0.672 |
| | Mean FD | -0.03 | 0.05 | -0.73 | 0.466 |
| | Random effects | σ | | | |
| | ID | 0.25 | | | |
| | Residual | 0.57 | | | |
| S01 | Predictors | β | SE | z | p |
| | (Intercept) | 0.02 | 0.08 | 0.27 | 0.785 |
| | Age | -0.04 | 0.07 | -0.65 | 0.520 |
| | Hemisphere (right) | 0.00 | 0.08 | 0.06 | 0.951 |
| | Mean FD | -0.06 | 0.08 | -0.78 | 0.435 |

| | | | | | |
|---|---|---|---|---|---|
| | Random effects | σ | | | |
| | ID | 0.56 | | | |
| | Residual | 0.59 | | | |
| | | ***MMPFC*** | | | |
| F01 | Predictors | β | SE | z | p |
| | (Intercept) | -0.07 | 0.08 | -0.91 | 0.365 |
| | Age | -0.03 | 0.07 | -0.44 | 0.662 |
| | Hemisphere (right) | -0.39 | 0.09 | -4.11 | **$7.941 \times 10^{-5}$** |
| | Mean FD | -0.12 | 0.07 | -1.58 | 0.117 |
| | Random effects | σ | | | |
| | ID | 0.49 | | | |
| | Residual | 0.68 | | | |
| F09 | Predictors | β | SE | z | p |
| | (Intercept) | -0.21 | 0.08 | -2.66 | 0.009 |
| | Age | -0.10 | 0.08 | -1.32 | 0.190 |
| | Hemisphere (right) | -0.30 | 0.07 | -4.49 | **$1.773 \times 10^{-5}$** |
| | Mean FD | -0.08 | 0.08 | -1.05 | 0.297 |
| | Random effects | σ | | | |
| | ID | 0.69 | | | |
| | Residual | 0.50 | | | |
| F11 | Predictors | β | SE | z | p |
| | (Intercept) | 0.00 | 0.07 | 0.03 | 0.977 |
| | Age | -0.11 | 0.06 | -1.73 | 0.086 |
| | Hemisphere (right) | -0.14 | 0.07 | -2.01 | 0.047 |
| | Mean FD | -0.10 | 0.06 | -1.57 | 0.119 |
| | Random effects | σ | | | |
| | ID | 0.54 | | | |
| | Residual | 0.50 | | | |
| F04 | Predictors | β | SE | z | p |
| | (Intercept) | -0.19 | 0.07 | -2.80 | 0.006 |
| | Age | 0.01 | 0.06 | 0.14 | 0.893 |
| | Hemisphere (right) | -0.01 | 0.05 | -0.27 | 0.788 |
| | Mean FD | -0.02 | 0.06 | -0.32 | 0.750 |
| | Random effects | σ | | | |
| | ID | 0.62 | | | |
| | Residual | 0.39 | | | |
| F10 | Predictors | β | SE | z | p |
| | (Intercept) | 0.01 | 0.06 | 0.26 | 0.794 |
| | Age | -0.04 | 0.06 | -0.73 | 0.468 |
| | Hemisphere (right) | 0.27 | 0.06 | 4.78 | **$5.300 \times 10^{-6}$** |
| | Mean FD | -0.01 | 0.05 | -0.15 | 0.880 |
| | Age x Hemisphere (right) | 0.15 | 0.06 | 2.60 | 0.011 |
| | Random effects | σ | | | |
| | ID | 0.40 | | | |
| | Residual | 0.43 | | | |
| F12 | Predictors | β | SE | z | p |
| | (Intercept) | 0.12 | 0.07 | 1.64 | 0.104 |
| | Age | -0.06 | 0.07 | -0.91 | 0.366 |
| | Hemisphere (right) | 0.18 | 0.06 | 3.13 | **0.002** |
| | Mean FD | 0.04 | 0.06 | 0.58 | 0.565 |
| | Age x Hemisphere (right) | 0.13 | 0.06 | 2.18 | 0.031 |
| | Random effects | σ | | | |
| | ID | 0.62 | | | |
| | Residual | 0.44 | | | |
| F07 | Predictors | β | SE | z | p |
| | (Intercept) | -0.04 | 0.07 | -0.62 | 0.538 |
| | Age | -0.04 | 0.07 | -0.58 | 0.562 |
| | Hemisphere (right) | -0.09 | 0.08 | -1.08 | 0.283 |
| | Mean FD | 0.01 | 0.06 | 0.23 | 0.818 |
| | Age x Hemisphere (right) | -0.19 | 0.08 | -2.35 | 0.021 |
| | Random effects | σ | | | |
| | ID | 0.44 | | | |
| | Residual | 0.59 | | | |

| | | | | | |
|---|---|---|---|---|---|
| F08 | Predictors | β | SE | z | p |
| | (Intercept) | 0.18 | 0.07 | 2.71 | 0.007 |
| | Age | 0.08 | 0.06 | 1.25 | 0.213 |
| | Hemisphere (right) | -0.02 | 0.07 | -0.30 | 0.768 |
| | Mean FD | 0.01 | 0.06 | 0.14 | 0.889 |
| | Random effects | σ | | | |
| | ID | 0.52 | | | |
| | Residual | 0.49 | | | |
| F06 | Predictors | β | SE | z | p |
| | (Intercept) | -0.14 | 0.07 | -2.22 | 0.028 |
| | Age | -0.01 | 0.06 | -0.27 | 0.789 |
| | Hemisphere (right) | -0.16 | 0.07 | -2.32 | 0.022 |
| | Mean FD | 0.06 | 0.06 | 0.96 | 0.338 |
| | Random effects | σ | | | |
| | ID | 0.47 | | | |
| | Residual | 0.50 | | | |
| F05 | Predictors | β | SE | z | p |
| | (Intercept) | -0.23 | 0.07 | -3.42 | **7.574 x 10$^{-4}$** |
| | Age | -0.23 | 0.06 | -3.61 | **3.798 x 10$^{-4}$** |
| | Hemisphere (right) | 0.05 | 0.09 | 0.61 | 0.545 |
| | Mean FD | 0.10 | 0.05 | 1.96 | 0.052 |
| | Age x hemisphere (right) | 0.19 | 0.08 | 2.28 | 0.025 |
| | Random effects | σ | | | |
| | ID | 0.26 | | | |
| | Residual | 0.64 | | | |
| F02 | Predictors | β | SE | z | p |
| | (Intercept) | 0.22 | 0.08 | 2.87 | 0.005 |
| | Age | 0.19 | 0.07 | 2.70 | 0.008 |
| | Hemisphere (right) | -0.15 | 0.06 | -2.54 | 0.013 |
| | Mean FD | -0.03 | 0.08 | -0.42 | 0.677 |
| | Random effects | σ | | | |
| | ID | 0.69 | | | |
| | Residual | 0.42 | | | |
| F03 | Predictors | β | SE | z | p |
| | (Intercept) | 0.16 | 0.08 | 2.16 | 0.033 |
| | Age | 0.11 | 0.07 | 1.69 | 0.095 |
| | Hemisphere (right) | 0.08 | 0.06 | 1.31 | 0.195 |
| | Mean FD | -0.16 | 0.08 | -1.95 | 0.054 |
| | Random effects | σ | | | |
| | ID | 0.64 | | | |
| | Residual | 0.44 | | | |
| S07 | Predictors | β | SE | z | p |
| | (Intercept) | -0.10 | 0.09 | -1.13 | 0.259 |
| | Age | -0.04 | 0.08 | -0.50 | 0.619 |
| | Hemisphere (right) | -0.42 | 0.08 | -5.21 | **9.598 x 10$^{-7}$** |
| | Mean FD | -0.18 | 0.09 | -1.90 | 0.060 |
| | Random effects | σ | | | |
| | ID | 0.73 | | | |
| | Residual | 0.58 | | | |
| S02 | Predictors | β | SE | z | p |
| | (Intercept) | -0.19 | 0.07 | -2.80 | 0.006 |
| | Age | 0.01 | 0.06 | 0.14 | 0.893 |
| | Hemisphere (right) | -0.01 | 0.05 | -0.27 | 0.788 |
| | Mean FD | -0.02 | 0.06 | -0.32 | 0.750 |
| | Random effects | σ | | | |
| | ID | 0.62 | | | |
| | Residual | 0.39 | | | |
| S03 | Predictors | β | SE | z | p |
| | (Intercept) | 0.12 | 0.06 | 2.04 | 0.043 |
| | Age | 0.07 | 0.05 | 1.29 | 0.201 |
| | Hemisphere (right) | 0.02 | 0.06 | 0.31 | 0.756 |
| | Mean FD | 0.02 | 0.05 | 0.41 | 0.682 |
| | Random effects | σ | | | |
| | ID | 0.45 | | | |
| | Residual | 0.43 | | | |

| | | | | | |
|---|---|---|---|---|---|
| S04 | Predictors | β | SE | z | p |
| | (Intercept) | 0.47 | 0.06 | 7.35 | **1.338 x $10^{-11}$** |
| | Age | 0.13 | 0.06 | 2.18 | 0.031 |
| | Hemisphere (right) | 0.12 | 0.05 | 2.37 | 0.019 |
| | Mean FD | -0.03 | 0.07 | -0.43 | 0.670 |
| | Random effects | σ | | | |
| | ID | 0.56 | | | |
| | Residual | 0.37 | | | |
| S12 | Predictors | β | SE | z | p |
| | (Intercept) | 0.26 | 0.07 | 3.83 | **1.800 x $10^{-4}$** |
| | Age | -0.01 | 0.06 | -0.14 | 0.889 |
| | Hemisphere (right) | 0.03 | 0.06 | 0.46 | 0.646 |
| | Mean FD | -0.03 | 0.06 | -0.49 | 0.627 |
| | Random effects | σ | | | |
| | ID | 0.52 | | | |
| | Residual | 0.47 | | | |
| S11 | Predictors | β | SE | z | p |
| | (Intercept) | 0.22 | 0.07 | 3.30 | **0.001** |
| | Age | -0.05 | 0.06 | -0.98 | 0.328 |
| | Hemisphere (right) | 0.03 | 0.07 | 0.34 | 0.733 |
| | Mean FD | -0.01 | 0.06 | -0.18 | 0.854 |
| | Random effects | σ | | | |
| | ID | 0.45 | | | |
| | Residual | 0.56 | | | |
| S10 | Predictors | β | SE | z | p |
| | (Intercept) | -0.19 | 0.07 | -2.72 | 0.007 |
| | Age | -0.02 | 0.06 | -0.41 | 0.682 |
| | Hemisphere (right) | 0.07 | 0.07 | 1.02 | 0.311 |
| | Mean FD | -0.04 | 0.06 | -0.65 | 0.519 |
| | Random effects | σ | | | |
| | ID | 0.48 | | | |
| | Residual | 0.54 | | | |
| S06 | Predictors | β | SE | z | p |
| | (Intercept) | -0.03 | 0.06 | -0.43 | 0.665 |
| | Age | -0.02 | 0.05 | -0.38 | 0.701 |
| | Hemisphere (right) | 0.07 | 0.06 | 1.17 | 0.246 |
| | Mean FD | 0.09 | 0.06 | 1.68 | 0.095 |
| | Random effects | σ | | | |
| | ID | 0.48 | | | |
| | Residual | 0.45 | | | |
| S08 | Predictors | β | SE | z | p |
| | (Intercept) | 0.02 | 0.07 | 0.29 | 0.771 |
| | Age | -0.07 | 0.06 | -1.24 | 0.218 |
| | Hemisphere (right) | -0.05 | 0.06 | -0.91 | 0.363 |
| | Mean FD | 0.03 | 0.07 | 0.51 | 0.611 |
| | Random effects | σ | | | |
| | ID | 0.52 | | | |
| | Residual | 0.44 | | | |
| S05 | Predictors | β | SE | z | p |
| | (Intercept) | 0.24 | 0.06 | 4.03 | **8.682 x $10^{-5}$** |
| | Age | 0.10 | 0.05 | 2.00 | 0.048 |
| | Hemisphere (right) | -0.03 | 0.05 | -0.60 | 0.551 |
| | Mean FD | -0.04 | 0.05 | -0.74 | 0.462 |
| | Random effects | σ | | | |
| | ID | 0.47 | | | |
| | Residual | 0.40 | | | |
| S09 | Predictors | β | SE | z | p |
| | (Intercept) | -0.04 | 0.06 | -0.62 | 0.538 |
| | Age | 0.02 | 0.04 | 0.36 | 0.718 |
| | Hemisphere (right) | -0.02 | 0.09 | -0.20 | 0.844 |
| | Mean FD | -0.04 | 0.05 | -0.85 | 0.395 |
| | Random effects | σ | | | |
| | ID | 0.13 | | | |
| | Residual | 0.65 | | | |

| | Predictors | β | SE | z | p |
|---|---|---|---|---|---|
| S01 | (Intercept) | 0.21 | 0.08 | 2.55 | 0.012 |
| | Age | 0.07 | 0.08 | 0.89 | 0.374 |
| | Hemisphere (right) | -0.01 | 0.06 | -0.11 | 0.912 |
| | Mean FD | -0.15 | 0.09 | -1.76 | 0.082 |
| | Random effects | σ | | | |
| | ID | 0.71 | | | |
| | Residual | 0.47 | | | |
| | ***Amygdala*** | | | | |
| F01 | Predictors | β | SE | z | p |
| | (Intercept) | -0.17 | 0.08 | -2.03 | 0.044 |
| | Age | -0.05 | 0.07 | -0.75 | 0.457 |
| | Hemisphere (right) | -0.19 | 0.09 | -1.99 | 0.050 |
| | Mean FD | 0.06 | 0.08 | 0.75 | 0.455 |
| | Random effects | σ | | | |
| | ID | 0.51 | | | |
| | Residual | 0.68 | | | |
| F09 | Predictors | β | SE | z | p |
| | (Intercept) | -0.27 | 0.08 | -3.39 | **8.830 x $10^{-4}$** |
| | Age | -0.12 | 0.07 | -1.68 | 0.096 |
| | Hemisphere (right) | -0.05 | 0.07 | -0.69 | 0.491 |
| | Mean FD | -0.04 | 0.08 | -0.56 | 0.577 |
| | Random effects | σ | | | |
| | ID | 0.62 | | | |
| | Residual | 0.55 | | | |
| F11 | Predictors | β | SE | z | p |
| | (Intercept) | -0.16 | 0.08 | -1.94 | 0.054 |
| | Age | -0.10 | 0.08 | -1.26 | 0.209 |
| | Hemisphere (right) | -0.08 | 0.07 | -1.19 | 0.238 |
| | Mean FD | -0.01 | 0.07 | -0.13 | 0.898 |
| | Random effects | σ | | | |
| | ID | 0.69 | | | |
| | Residual | 0.48 | | | |
| F04 | Predictors | β | SE | z | p |
| | (Intercept) | -0.15 | 0.06 | -2.55 | 0.012 |
| | Age | -0.16 | 0.06 | -2.69 | 0.008 |
| | Hemisphere (right) | 0.07 | 0.07 | 0.90 | 0.369 |
| | Mean FD | -0.03 | 0.05 | -0.6 | 0.547 |
| | Age x hemisphere (right) | 0.18 | 0.07 | 2.50 | 0.014 |
| | Random effects | σ | | | |
| | ID | 0.34 | | | |
| | Residual | 0.54 | | | |
| F10 | Predictors | β | SE | z | p |
| | (Intercept) | 0.28 | 0.05 | 5.30 | **3.098 x $10^{-7}$** |
| | Age | -0.05 | 0.04 | -1.21 | 0.229 |
| | Hemisphere (right) | -0.03 | 0.06 | -0.51 | 0.609 |
| | Mean FD | -0.03 | 0.04 | -0.57 | 0.572 |
| | Random effects | σ | | | |
| | ID | 0.35 | | | |
| | Residual | 0.46 | | | |
| F12 | Predictors | β | SE | z | p |
| | (Intercept) | 0.17 | 0.05 | 3.18 | **0.002** |
| | Age | -0.05 | 0.05 | -1.03 | 0.305 |
| | Hemisphere (right) | 0.04 | 0.05 | 0.82 | 0.412 |
| | Mean FD | -0.08 | 0.05 | -1.84 | 0.069 |
| | Random effects | σ | | | |
| | ID | 0.42 | | | |
| | Residual | 0.36 | | | |
| F07 | Predictors | β | SE | z | p |
| | (Intercept) | 0.14 | 0.06 | 2.51 | 0.013 |
| | Age | -0.01 | 0.05 | -0.27 | 0.786 |
| | Hemisphere (right) | -0.08 | 0.05 | -1.46 | 0.147 |
| | Mean FD | -0.05 | 0.05 | -0.98 | 0.331 |
| | Random effects | σ | | | |

| | | | | | |
|---|---|---|---|---|---|
| | ID | 0.43 | | | |
| | Residual | 0.40 | | | |
| F08 | Predictors | β | SE | z | p |
| | (Intercept) | -0.01 | 0.05 | -0.14 | 0.886 |
| | Age | -0.08 | 0.05 | -1.55 | 0.123 |
| | Hemisphere (right) | 0.07 | 0.05 | 1.52 | 0.132 |
| | Mean FD | -0.02 | 0.05 | -0.46 | 0.646 |
| | Random effects | σ | | | |
| | ID | 0.44 | | | |
| | Residual | 0.37 | | | |
| F06 | Predictors | β | SE | z | p |
| | (Intercept) | 0.00 | 0.05 | 0.01 | 0.990 |
| | Age | 0.03 | 0.05 | 0.67 | 0.503 |
| | Hemisphere (right) | -0.03 | 0.05 | -0.56 | 0.576 |
| | Mean FD | 0.05 | 0.05 | 0.96 | 0.341 |
| | Random effects | σ | | | |
| | ID | 0.41 | | | |
| | Residual | 0.36 | | | |
| F05 | Predictors | β | SE | z | p |
| | (Intercept) | 0.08 | 0.05 | 1.49 | 0.137 |
| | Age | -0.01 | 0.04 | -0.28 | 0.778 |
| | Hemisphere (right) | 0.15 | 0.07 | 2.20 | 0.030 |
| | Mean FD | -0.02 | 0.04 | -0.47 | 0.637 |
| | Random effects | σ | | | |
| | ID | 0.27 | | | |
| | Residual | 0.49 | | | |
| F02 | Predictors | β | SE | z | p |
| | (Intercept) | 0.11 | 0.07 | 1.62 | 0.108 |
| | Age | -0.01 | 0.06 | -0.09 | 0.932 |
| | Hemisphere (right) | 0.05 | 0.05 | 0.98 | 0.328 |
| | Mean FD | -0.04 | 0.07 | -0.57 | 0.571 |
| | Random effects | σ | | | |
| | ID | 0.60 | | | |
| | Residual | 0.35 | | | |
| F03 | Predictors | β | SE | z | p |
| | (Intercept) | -0.03 | 0.07 | -0.42 | 0.675 |
| | Age | 0.04 | 0.06 | 0.75 | 0.454 |
| | Hemisphere (right) | 0.23 | 0.07 | 3.23 | **0.002** |
| | Mean FD | 0.05 | 0.06 | 0.90 | 0.371 |
| | Random effects | σ | | | |
| | ID | 0.48 | | | |
| | Residual | 0.53 | | | |
| S07 | Predictors | β | SE | z | p |
| | (Intercept) | -0.2 | 0.08 | -2.37 | 0.019 |
| | Age | -0.11 | 0.07 | -1.51 | 0.133 |
| | Hemisphere (right) | -0.17 | 0.07 | -2.35 | 0.021 |
| | Mean FD | -0.07 | 0.09 | -0.85 | 0.395 |
| | Random effects | σ | | | |
| | ID | 0.68 | | | |
| | Residual | 0.51 | | | |
| S02 | Predictors | β | SE | z | p |
| | (Intercept) | -0.01 | 0.06 | -0.16 | 0.873 |
| | Age | 0.01 | 0.05 | 0.29 | 0.773 |
| | Hemisphere (right) | -0.07 | 0.07 | -1.09 | 0.277 |
| | Mean FD | 0.03 | 0.05 | 0.66 | 0.509 |
| | Random effects | σ | | | |
| | ID | 0.36 | | | |
| | Residual | 0.51 | | | |
| S03 | Predictors | β | SE | z | p |
| | (Intercept) | 0.14 | 0.06 | 2.52 | 0.013 |
| | Age | -0.02 | 0.05 | -0.36 | 0.720 |
| | Hemisphere (right) | -0.05 | 0.06 | -0.83 | 0.407 |
| | Mean FD | -0.06 | 0.05 | -1.29 | 0.200 |
| | Random effects | σ | | | |

| | | | | | |
|---|---|---|---|---|---|
| | ID | 0.36 | | | |
| | Residual | 0.46 | | | |
| S04 | Predictors | β | SE | z | p |
| | (Intercept) | 0.12 | 0.06 | 2.10 | 0.038 |
| | Age | -0.01 | 0.05 | -0.27 | 0.790 |
| | Hemisphere (right) | 0.05 | 0.05 | 1.02 | 0.311 |
| | Mean FD | -0.02 | 0.06 | -0.43 | 0.665 |
| | Random effects | σ | | | |
| | ID | 0.44 | | | |
| | Residual | 0.34 | | | |
| S12 | Predictors | β | SE | z | p |
| | (Intercept) | 0.17 | 0.05 | 3.18 | **0.002** |
| | Age | -0.05 | 0.05 | -1.03 | 0.305 |
| | Hemisphere (right) | 0.04 | 0.05 | 0.82 | 0.412 |
| | Mean FD | -0.08 | 0.05 | -1.84 | 0.069 |
| | Random effects | σ | | | |
| | ID | 0.42 | | | |
| | Residual | 0.36 | | | |
| S11 | Predictors | β | SE | z | p |
| | (Intercept) | 0.02 | 0.06 | 0.30 | 0.763 |
| | Age | -0.06 | 0.05 | -1.14 | 0.255 |
| | Hemisphere (right) | 0.09 | 0.05 | 1.74 | 0.084 |
| | Mean FD | 0.01 | 0.05 | 0.19 | 0.849 |
| | Random effects | σ | | | |
| | ID | 0.50 | | | |
| | Residual | 0.37 | | | |
| S10 | Predictors | β | SE | z | p |
| | (Intercept) | -0.07 | 0.06 | -1.17 | 0.242 |
| | Age | 0.00 | 0.06 | -0.03 | 0.976 |
| | Hemisphere (right) | 0.05 | 0.05 | 0.95 | 0.344 |
| | Mean FD | -0.03 | 0.06 | -0.46 | 0.648 |
| | Random effects | σ | | | |
| | ID | 0.54 | | | |
| | Residual | 0.36 | | | |
| S06 | Predictors | β | SE | z | p |
| | (Intercept) | -0.14 | 0.05 | -2.69 | 0.008 |
| | Age | 0.03 | 0.05 | 0.67 | 0.507 |
| | Hemisphere (right) | 0.10 | 0.05 | 2.28 | 0.024 |
| | Mean FD | 0.07 | 0.05 | 1.53 | 0.129 |
| | Random effects | σ | | | |
| | ID | 0.43 | | | |
| | Residual | 0.34 | | | |
| S08 | Predictors | β | SE | z | p |
| | (Intercept) | 0.03 | 0.06 | 0.48 | 0.634 |
| | Age | -0.03 | 0.05 | -0.69 | 0.494 |
| | Hemisphere (right) | 0.06 | 0.06 | 0.97 | 0.335 |
| | Mean FD | -0.01 | 0.05 | -0.13 | 0.899 |
| | Random effects | σ | | | |
| | ID | 0.43 | | | |
| | Residual | 0.43 | | | |
| S05 | Predictors | β | SE | z | p |
| | (Intercept) | -0.14 | 0.07 | -2.02 | 0.045 |
| | Age | 0.12 | 0.06 | 2.00 | 0.048 |
| | Hemisphere (right) | 0.02 | 0.05 | 0.32 | 0.747 |
| | Mean FD | 0.02 | 0.07 | 0.28 | 0.78 |
| | Random effects | σ | | | |
| | ID | 0.59 | | | |
| | Residual | 0.40 | | | |
| S09 | Predictors | β | SE | z | p |
| | (Intercept) | 0.18 | 0.06 | 3.29 | **0.001** |
| | Age | 0.08 | 0.05 | 1.57 | 0.120 |
| | Hemisphere (right) | 0.02 | 0.05 | 0.39 | 0.696 |
| | Mean FD | 0.07 | 0.05 | 1.38 | 0.172 |
| | Random effects | σ | | | |

| | | | | | |
|---|---|---|---|---|---|
| | ID | 0.46 | | | |
| | Residual | 0.36 | | | |
| S01 | Predictors | β | SE | z | p |
| | (Intercept) | 0.09 | 0.09 | 1.00 | 0.319 |
| | Age | 0.04 | 0.08 | 0.44 | 0.664 |
| | Hemisphere (right) | 0.03 | 0.06 | 0.45 | 0.656 |
| | Mean FD | 0.08 | 0.11 | 0.79 | 0.431 |
| | Random effects | σ | | | |
| | ID | 0.77 | | | |
| | Residual | 0.41 | | | |
| | | ***STS*** | | | |
| F01 | Predictors | β | SE | z | p |
| | (Intercept) | 0.00 | 0.08 | 0.03 | 0.974 |
| | Age | -0.13 | 0.06 | -2.22 | 0.029 |
| | Hemisphere (right) | -0.28 | 0.10 | -2.81 | 0.006 |
| | Mean FD | -0.07 | 0.06 | -1.02 | 0.308 |
| | Random effects | σ | | | |
| | ID | 0.30 | | | |
| | Residual | 0.73 | | | |
| F09 | Predictors | β | SE | z | p |
| | (Intercept) | 0.09 | 0.07 | 1.35 | 0.180 |
| | Age | 0.05 | 0.07 | 0.64 | 0.525 |
| | Hemisphere (right) | -0.59 | 0.09 | -6.64 | **$1.228 \times 10^{-9}$** |
| | Mean FD | -0.11 | 0.06 | -1.96 | 0.052 |
| | Age x hemisphere (right) | -0.24 | 0.09 | -2.62 | 0.010 |
| | Random effects | σ | | | |
| | ID | 0.31 | | | |
| | Residual | 0.67 | | | |
| F11 | Predictors | β | SE | z | p |
| | (Intercept) | -0.26 | 0.07 | -3.93 | **$1.140 \times 10^{-4}$** |
| | Age | -0.10 | 0.05 | -2.13 | 0.036 |
| | Hemisphere (right) | -0.04 | 0.09 | -0.39 | 0.699 |
| | Mean FD | 0.08 | 0.05 | 1.82 | 0.071 |
| | Random effects | σ | | | |
| | ID | 0.08 | | | |
| | Residual | 0.67 | | | |
| F04 | Predictors | β | SE | z | p |
| | (Intercept) | -0.33 | 0.05 | -5.94 | **$1.131 \times 10^{-8}$** |
| | Age | 0.04 | 0.04 | 0.88 | 0.379 |
| | Hemisphere (right) | 0.03 | 0.07 | 0.38 | 0.705 |
| | Mean FD | -0.04 | 0.04 | -1.07 | 0.288 |
| | Random effects | σ | | | |
| | ID | 0.16 | | | |
| | Residual | 0.55 | | | |
| F10 | Predictors | β | SE | z | p |
| | (Intercept) | 0.34 | 0.06 | 5.62 | **$5.497 \times 10^{-8}$** |
| | Age | 0.14 | 0.04 | 3.42 | **$7.311 \times 10^{-4}$** |
| | Hemisphere (right) | -0.01 | 0.08 | -0.11 | 0.912 |
| | Mean FD | -0.02 | 0.04 | -0.55 | 0.583 |
| | Random effects | σ | | | |
| | ID | 0.00 | | | |
| | Residual | 0.64 | | | |
| F12 | Predictors | β | SE | z | p |
| | (Intercept) | 0.17 | 0.06 | 2.78 | 0.006 |
| | Age | 0.07 | 0.05 | 1.52 | 0.132 |
| | Hemisphere (right) | 0.06 | 0.08 | 0.71 | 0.478 |
| | Mean FD | -0.04 | 0.05 | -0.77 | 0.444 |
| | Random effects | σ | | | |
| | ID | 0.24 | | | |
| | Residual | 0.60 | | | |
| F07 | Predictors | β | SE | z | p |
| | (Intercept) | 0.17 | 0.06 | 2.71 | 0.007 |
| | Age | 0.03 | 0.05 | 0.59 | 0.559 |
| | Hemisphere (right) | -0.36 | 0.07 | -4.86 | **$3.954 \times 10^{-6}$** |

| | | | | | |
|---|---|---|---|---|---|
| | Mean FD | 0.00 | 0.05 | 0.09 | 0.932 |
| | Random effects | σ | | | |
| | ID | 0.35 | | | |
| | Residual | 0.55 | | | |
| F08 | Predictors | β | SE | z | p |
| | (Intercept) | 0.32 | 0.06 | 4.98 | **1.291 x $10^{-6}$** |
| | Age | 0.11 | 0.05 | 2.37 | 0.020 |
| | Hemisphere (right) | -0.24 | 0.09 | -2.73 | 0.007 |
| | Mean FD | 0.00 | 0.05 | 0.08 | 0.933 |
| | Random effects | σ | | | |
| | ID | 0.20 | | | |
| | Residual | 0.64 | | | |
| F06 | Predictors | β | SE | z | p |
| | (Intercept) | -0.10 | 0.06 | -1.79 | 0.075 |
| | Age | -0.02 | 0.05 | -0.53 | 0.600 |
| | Hemisphere (right) | -0.02 | 0.07 | -0.33 | 0.743 |
| | Mean FD | 0.04 | 0.05 | 0.80 | 0.424 |
| | Random effects | σ | | | |
| | ID | 0.31 | | | |
| | Residual | 0.52 | | | |
| F05 | Predictors | β | SE | z | p |
| | (Intercept) | -0.21 | 0.07 | -3.07 | **0.002** |
| | Age | 0.04 | 0.05 | 0.88 | 0.382 |
| | Hemisphere (right) | 0.23 | 0.10 | 2.44 | 0.015 |
| | Mean FD | 0.05 | 0.05 | 0.99 | 0.324 |
| | Random effects | σ | | | |
| | ID | 0.00 | | | |
| | Residual | 0.72 | | | |
| F02 | Predictors | β | SE | z | p |
| | (Intercept) | 0.12 | 0.06 | 1.99 | 0.048 |
| | Age | 0.07 | 0.05 | 1.44 | 0.152 |
| | Hemisphere (right) | 0.07 | 0.08 | 0.91 | 0.365 |
| | Mean FD | -0.03 | 0.05 | -0.62 | 0.537 |
| | Random effects | σ | | | |
| | ID | 0.22 | | | |
| | Residual | 0.60 | | | |
| F03 | Predictors | β | SE | z | p |
| | (Intercept) | 0.20 | 0.06 | 3.10 | **0.002** |
| | Age | -0.01 | 0.05 | -0.13 | 0.893 |
| | Hemisphere (right) | -0.03 | 0.08 | -0.31 | 0.756 |
| | Mean FD | -0.07 | 0.06 | -1.20 | 0.233 |
| | Random effects | σ | | | |
| | ID | 0.28 | | | |
| | Residual | 0.61 | | | |
| S07 | Predictors | β | SE | z | p |
| | (Intercept) | 0.00 | 0.08 | 0.03 | 0.974 |
| | Age | -0.13 | 0.06 | -2.22 | 0.029 |
| | Hemisphere (right) | -0.28 | 0.10 | -2.81 | 0.006 |
| | Mean FD | -0.07 | 0.06 | -1.02 | 0.308 |
| | Random effects | σ | | | |
| | ID | 0.30 | | | |
| | Residual | 0.73 | | | |
| S02 | Predictors | β | SE | z | p |
| | (Intercept) | -0.06 | 0.06 | -0.97 | 0.335 |
| | Age | -0.13 | 0.06 | -2.13 | 0.034 |
| | Hemisphere (right) | -0.24 | 0.08 | -2.82 | 0.006 |
| | Mean FD | 0.07 | 0.04 | 1.74 | 0.085 |
| | Age x Hemisphere (right) | 0.20 | 0.08 | 2.35 | 0.021 |
| | Random effects | σ | | | |
| | ID | 0.12 | | | |
| | Residual | 0.63 | | | |
| S03 | Predictors | β | SE | z | p |
| | (Intercept) | 0.39 | 0.05 | 7.62 | **7.641 x $10^{-13}$** |
| | Age | 0.05 | 0.04 | 1.36 | 0.176 |

| | | | | | |
|---|---|---|---|---|---|
| | Hemisphere (right) | 0.03 | 0.07 | 0.47 | 0.637 |
| | Mean FD | 0.08 | 0.04 | 1.87 | 0.064 |
| | Random effects | σ | | | |
| | ID | 0.18 | | | |
| | Residual | 0.51 | | | |
| S04 | Predictors | β | SE | z | p |
| | (Intercept) | 0.24 | 0.06 | 4.13 | **$5.466 \times 10^{-5}$** |
| | Age | 0.02 | 0.05 | 0.35 | 0.727 |
| | Hemisphere (right) | 0.07 | 0.07 | 0.88 | 0.382 |
| | Mean FD | 0.05 | 0.05 | 1.02 | 0.311 |
| | Random effects | σ | | | |
| | ID | 0.27 | | | |
| | Residual | 0.53 | | | |
| S12 | Predictors | β | SE | z | p |
| | (Intercept) | 0.17 | 0.06 | 2.78 | 0.006 |
| | Age | 0.07 | 0.05 | 1.52 | 0.132 |
| | Hemisphere (right) | 0.06 | 0.08 | 0.71 | 0.478 |
| | Mean FD | -0.04 | 0.05 | -0.77 | 0.444 |
| | Random effects | σ | | | |
| | ID | 0.24 | | | |
| | Residual | 0.60 | | | |
| S11 | Predictors | β | SE | z | p |
| | (Intercept) | 0.00 | 0.07 | 0.04 | 0.972 |
| | Age | -0.01 | 0.05 | -0.21 | 0.837 |
| | Hemisphere (right) | 0.32 | 0.08 | 3.76 | **$2.696 \times 10^{-6}$** |
| | Mean FD | -0.05 | 0.05 | -0.97 | 0.335 |
| | Random effects | σ | | | |
| | ID | 0.32 | | | |
| | Residual | 0.63 | | | |
| S10 | Predictors | β | SE | z | p |
| | (Intercept) | -0.12 | 0.06 | -1.93 | 0.055 |
| | Age | 0.01 | 0.04 | 0.33 | 0.739 |
| | Hemisphere (right) | 0.00 | 0.09 | -0.03 | 0.980 |
| | Mean FD | -0.04 | 0.04 | -0.79 | 0.428 |
| | Random effects | σ | | | |
| | ID | 0.00 | | | |
| | Residual | 0.64 | | | |
| S06 | Predictors | β | SE | z | p |
| | (Intercept) | -0.21 | 0.07 | -3.07 | **0.002** |
| | Age | 0.04 | 0.05 | 0.88 | 0.382 |
| | Hemisphere (right) | 0.23 | 0.10 | 2.44 | 0.015 |
| | Mean FD | 0.05 | 0.05 | 0.99 | 0.324 |
| | Random effects | σ | | | |
| | ID | 0.00 | | | |
| | Residual | 0.72 | | | |
| S08 | Predictors | β | SE | z | p |
| | (Intercept) | 0.09 | 0.05 | 1.63 | 0.104 |
| | Age | 0.01 | 0.05 | 0.14 | 0.887 |
| | Hemisphere (right) | -0.02 | 0.07 | -0.28 | 0.782 |
| | Mean FD | 0.00 | 0.05 | -0.10 | 0.921 |
| | Age x hemisphere (right) | -0.13 | 0.06 | -2.02 | 0.046 |
| | Random effects | σ | | | |
| | ID | 0.28 | | | |
| | Residual | 0.48 | | | |
| S05 | Predictors | β | SE | z | p |
| | (Intercept) | -0.03 | 0.06 | -0.47 | 0.640 |
| | Age | 0.01 | 0.05 | 0.16 | 0.876 |
| | Hemisphere (right) | -0.02 | 0.08 | -0.25 | 0.807 |
| | Mean FD | 0.01 | 0.05 | 0.13 | 0.901 |
| | Random effects | σ | | | |
| | ID | 0.28 | | | |
| | Residual | 0.57 | | | |
| S09 | Predictors | β | SE | z | p |
| | (Intercept) | -0.20 | 0.05 | -3.60 | **$3.944 \times 10^{-4}$** |

| | | | | | |
|---|---|---|---|---|---|
| | Age | -0.03 | 0.04 | -0.60 | 0.549 |
| | Hemisphere (right) | 0.27 | 0.07 | 3.97 | **$1.255 \times 10^{-4}$** |
| | Mean FD | 0.01 | 0.04 | 0.28 | 0.782 |
| | Random effects | σ | | | |
| | ID | 0.25 | | | |
| | Residual | 0.52 | | | |
| S01 | Predictors | β | SE | z | P |
| | (Intercept) | 0.30 | 0.07 | 4.10 | **$5.976 \times 10^{-5}$** |
| | Age | 0.03 | 0.06 | 0.55 | 0.584 |
| | Hemisphere (right) | -0.06 | 0.09 | -0.66 | 0.509 |
| | Mean FD | -0.05 | 0.07 | -0.70 | 0.485 |
| | Random effects | σ | | | |
| | ID | 0.34 | | | |
| | Residual | 0.69 | | | |

*Face events are denoted F01-F12, scene events are denoted S01-S12. Events are listed in order of appearance in the movie (see Figure 1 in the main text), with event labels reflecting the ranking of average magnitude of response of peak timepoints in adults, as reported previously (Kamps, Richardson et al., 2022). Interaction terms are denoted with "x"; age-by-hemisphere interaction terms that were not statistically significant were excluded from the final models. Reported βs are standardised, ps<0.004 are shown in bold for face and scene events (Bonferroni correction: p=0.05/12, for 12 events per ROI) and scientific notation is used for ps<0.001. N=117. FFA = Fusiform Face Area; MMPFC = Middle Medial Prefrontal Cortex; STS = Superior Temporal Sulcus; FD = Framewise Displacement; SE = Standard Error.*

***Table S4:*** *Age effects on functional connectivity between FFA, MMPFC, amygdala and STS*

| | Predictors | β | SE | z | p |
|---|---|---|---|---|---|
| ***Primary models*** *i.e., functional connectivity of right FFA as dependent variable* | | | | | |
| | Predictors | β | SE | z | p |
| Fx left FFA - right FFA | (Intercept) | -0.07 | 0.10 | -0.69 | 0.491 |
| | Age | 0.13 | 0.10 | 1.31 | 0.193 |
| | Mean FD | -0.15 | 0.10 | -1.55 | 0.124 |
| Fx right FFA - left/right MMPFC | (Intercept) | 0.11 | 0.09 | 1.23 | 0.221 |
| | Age | -0.01 | 0.09 | -0.16 | 0.873 |
| | Hemisphere (right) | -0.09 | 0.07 | -1.35 | 0.179 |
| | Mean FD | 0.14 | 0.09 | 1.56 | 0.121 |
| | Age x Hemisphere (right) | 0.16 | 0.07 | 2.44 | **0.016** |
| Fx right FFA - left/right amygdala | (Intercept) | -0.01 | 0.10 | -0.07 | 0.941 |
| | Age | 0.09 | 0.09 | 0.97 | 0.334 |
| | Hemisphere (right) | 0.01 | 0.08 | 0.11 | 0.909 |
| | Mean FD | 0.03 | 0.09 | 0.33 | 0.744 |
| Fx right FFA - left/right STS | (Intercept) | -0.01 | 0.09 | -0.14 | 0.891 |
| | Age | 0.05 | 0.07 | 0.68 | 0.499 |
| | Hemisphere (right) | -0.01 | 0.12 | -0.08 | 0.940 |
| | Mean FD | 0.15 | 0.07 | 2.20 | **0.030** |
| ***Secondary models*** *i.e., functional connectivity of left FFA, MMPFC, amygdala and STS as dependent variables* | | | | | |
| | Predictors | β | SE | z | p |
| Fx left FFA - left/right MMPFC | (Intercept) | 0.13 | 0.09 | 1.44 | 0.152 |
| | Age | 0.13 | 0.09 | 1.54 | 0.127 |
| | Hemisphere (right) | -0.11 | 0.08 | -1.45 | 0.148 |
| | Mean FD | 0.08 | 0.09 | 0.99 | 0.327 |
| Fx left FFA - left/right amygdala | (Intercept) | 0.06 | 0.10 | 0.62 | 0.536 |
| | Age | 0.03 | 0.09 | 0.38 | 0.708 |
| | Hemisphere (right) | -0.07 | 0.07 | -0.99 | 0.326 |
| | Mean FD | -0.05 | 0.09 | -0.61 | 0.541 |
| Fx left FFA - left/right STS | (Intercept) | 0.10 | 0.09 | 1.09 | 0.276 |
| | Age | -0.03 | 0.07 | -0.46 | 0.648 |
| | Hemisphere (right) | -0.12 | 0.13 | -0.96 | 0.340 |
| | Mean FD | 0.07 | 0.07 | 1.08 | 0.284 |
| Fx left - right MMPFC | (Intercept) | 0.13 | 0.09 | 1.40 | 0.163 |
| | Age | 0.00 | 0.09 | 0.01 | 0.994 |
| | Mean FD | 0.11 | 0.09 | 1.26 | 0.209 |
| Fx left MMPFC - left/right amygdala | (Intercept) | 0.10 | 0.09 | 1.06 | 0.290 |
| | Age | 0.02 | 0.09 | 0.20 | 0.840 |
| | Hemisphere (right) | 0.00 | 0.07 | -0.01 | 0.991 |
| | Mean FD | 0.08 | 0.09 | 0.88 | 0.380 |
| Fx left MMPFC - left/right STS | (Intercept) | 0.13 | 0.09 | 1.51 | 0.133 |
| | Age | 0.01 | 0.07 | 0.15 | 0.884 |
| | Hemisphere (right) | -0.04 | 0.11 | -0.39 | 0.696 |
| | Mean FD | 0.04 | 0.07 | 0.53 | 0.597 |
| Fx right MMPFC - left/right amygdala | (Intercept) | 0.05 | 0.09 | 0.56 | 0.576 |
| | Age | 0.10 | 0.09 | 1.11 | 0.268 |
| | Hemisphere (right) | 0.02 | 0.06 | 0.38 | 0.705 |
| | Mean FD | -0.11 | 0.09 | -1.26 | 0.211 |
| Fx right MMPFC - left/right STS | (Intercept) | 0.05 | 0.09 | 0.57 | 0.570 |
| | Age | 0.15 | 0.07 | 2.10 | **0.038** |
| | Hemisphere (right) | -0.11 | 0.12 | -0.87 | 0.384 |
| | Mean FD | 0.01 | 0.07 | 0.12 | 0.904 |
| Fx left - right amygdala | (Intercept) | -0.05 | 0.09 | -0.52 | 0.603 |
| | Age | -0.04 | 0.09 | -0.48 | 0.635 |
| | Mean FD | -0.28 | 0.09 | -3.10 | **0.002** |
| Fx left amygdala - left/right STS | (Intercept) | 0.02 | 0.09 | 0.27 | 0.790 |
| | Age | 0.05 | 0.07 | 0.70 | 0.483 |
| | Hemisphere (right) | -0.02 | 0.10 | -0.24 | 0.814 |
| | Mean FD | 0.03 | 0.07 | 0.38 | 0.703 |
| Fx right amygdala - left/right STS | (Intercept) | -0.05 | 0.09 | -0.58 | 0.564 |
| | Age | 0.00 | 0.07 | -0.03 | 0.978 |
| | Hemisphere (right) | 0.06 | 0.11 | 0.50 | 0.619 |
| | Mean FD | 0.04 | 0.07 | 0.54 | 0.588 |

| | | | | | |
|---|---|---|---|---|---|
| | (Intercept) | -0.10 | 0.09 | -1.10 | 0.272 |
| Fx left - right STS | Age | -0.03 | 0.09 | -0.30 | 0.763 |
| | Mean FD | -0.13 | 0.09 | -1.44 | 0.152 |

*Interaction terms are denoted with "x"; age-by-hemisphere interaction terms that were not statistically significant were excluded from the final models. Reported βs are standardised, ps<0.05 are shown in bold. N=117. FFA = Fusiform Face Area; MMPFC = Middle Medial Prefrontal Cortex; STS = Superior Temporal Sulcus; Fx = Functional Connectivity; FD = Framewise Displacement; SE = Standard Error.*

***Table S5:*** *Associations between functional maturity of left FFA and its functional connectivity with MMPFC, amygdala and STS*

| ***Fx left FFA - MMPFC*** | | | | |
|---|---|---|---|---|
| Predictors | β | SE | z | p |
| (Intercept) | 0.13 | 0.09 | 1.44 | 0.151 |
| Fm left FFA | -0.12 | 0.09 | -1.34 | 0.183 |
| MMPFC hemisphere (right) | -0.11 | 0.08 | -1.45 | 0.148 |
| Age | 0.16 | 0.09 | 1.86 | 0.066 |
| Mean FD | 0.07 | 0.09 | 0.85 | 0.395 |
| Random effects | σ | | | |
| ID | 0.81 | | | |
| Residual | 0.60 | | | |
| ***Fx left FFA - amygdala*** | | | | |
| Predictors | β | SE | z | p |
| (Intercept) | 0.06 | 0.10 | 0.62 | 0.536 |
| Fm left FFA | -0.09 | 0.09 | -0.92 | 0.357 |
| Amygdala hemisphere (right) | -0.07 | 0.07 | -0.99 | 0.326 |
| Age | 0.06 | 0.09 | 0.62 | 0.535 |
| Mean FD | -0.06 | 0.09 | -0.70 | 0.485 |
| Random effects | σ | | | |
| ID (σ) | 0.86 | | | |
| Residual (σ) | 0.57 | | | |
| ***Fx left FFA - STS*** | | | | |
| Predictors | β | SE | z | p |
| (Intercept) | 0.10 | 0.09 | 1.10 | 0.273 |
| Fm left FFA | -0.14 | 0.07 | -2.02 | **0.046** |
| STS hemisphere (right) | -0.12 | 0.13 | -0.96 | 0.340 |
| Age | 0.01 | 0.07 | 0.13 | 0.898 |
| Mean FD | 0.06 | 0.07 | 0.89 | 0.376 |
| Random effects | σ | | | |
| ID (σ) | 0.11 | | | |
| Residual (σ) | 0.98 | | | |

*Interaction terms of age-by-hemisphere-by-functional maturity (and all embedded two-way interaction terms) for each test were not statistically significant and were excluded from the final models. Reported βs are standardised, ps<0.05 are shown in bold. N=117. FFA = Fusiform Face Area; MMPFC = Middle Medial Prefrontal Cortex; STS = Superior Temporal Sulcus; Fm = Functional Maturity; Fx = Functional Connectivity; FD = Framewise Displacement; SE = Standard Error.*

***Table S6:*** *Associations between functional maturity of left FFA and functional maturity of MMPFC, amygdala and STS*

| | | | | |
|---|---|---|---|---|
| | | ***Fm MMPFC*** | | |
| Predictors | β | SE | z | p |
| (Intercept) | 0.00 | 0.08 | 0.00 | 1.000 |
| Fm left FFA | 0.11 | 0.08 | 1.48 | 0.143 |
| MMPFC hemisphere (right) | 0.00 | 0.09 | 0.00 | 1.000 |
| Age | 0.32 | 0.08 | 4.21 | **$5.190 \times 10^{-5}$** |
| Mean FD | -0.16 | 0.07 | -2.23 | **0.027** |
| Random effects | σ | | | |
| ID | 0.62 | | | |
| Residual | 0.68 | | | |
| | | ***Fm amygdala*** | | |
| Predictors | β | SE | z | p |
| (Intercept) | 0.00 | 0.09 | 0.00 | 1.000 |
| Fm left FFA | -0.13 | 0.09 | -1.53 | 0.130 |
| Amygdala hemisphere (right) | 0.00 | 0.08 | 0.00 | 1.000 |
| Age | 0.24 | 0.08 | 2.79 | **0.006** |
| Mean FD | 0.01 | 0.08 | 0.09 | 0.929 |
| Random effects | σ | | | |
| ID (σ) | 0.75 | | | |
| Residual (σ) | 0.64 | | | |
| | | ***Fm STS*** | | |
| Predictors | β | SE | z | p |
| (Intercept) | 0.00 | 0.09 | 0.00 | 1.000 |
| Fm left FFA | 0.16 | 0.07 | 2.38 | **0.019** |
| STS hemisphere (right) | 0.00 | 0.12 | 0.00 | 1.000 |
| Age | 0.20 | 0.07 | 2.90 | **0.005** |
| Mean FD | -0.09 | 0.06 | -1.46 | 0.146 |
| Random effects | σ | | | |
| ID (σ) | 0.22 | | | |
| Residual (σ) | 0.93 | | | |

*Interaction terms of age-by-hemisphere-by-functional maturity (and all embedded two-way interaction terms) for each test were not statistically significant and were excluded from the final models. Reported βs are standardised, ps<0.05 are shown in bold. N=117. FFA = Fusiform Face Area; MMPFC = Middle Medial Prefrontal Cortex; STS = Superior Temporal Sulcus; Fm = Functional Maturity; FD = Framewise Displacement; SE = Standard Error.*

***Table S7:*** *Age effects on FFA response lateralisation*

| | ***p<0.10*** | | | |
|---|---|---|---|---|
| Predictors | β | SE | z | p |
| (Intercept) | 0.00 | 0.09 | -0.03 | 0.976 |
| Age | -0.16 | 0.10 | -1.68 | 0.096 |
| Mean FD | 0.05 | 0.10 | 0.48 | 0.630 |

*Reported βs are standardised. N=110. FD = Framewise Displacement. See Appendix S3 for further results and discussion of lateralisation index metric.*

***Table S8:*** *Associations between lateralisation of FFA response and functional connectivity between right FFA, MMPFC, amygdala and STS*

| | | | | |
|---|---|---|---|---|
| ***Fx right FFA - MMPFC*** | | | | |
| Predictors | β | SE | z | p |
| (Intercept) | 0.10 | 0.10 | 1.06 | 0.290 |
| LI | 0.22 | 0.10 | 2.23 | **0.027** |
| MMPFC hemisphere (right) | -0.08 | 0.07 | -1.18 | 0.242 |
| Age | 0.08 | 0.09 | 0.86 | 0.392 |
| Mean FD | 0.12 | 0.09 | 1.29 | 0.201 |
| LI x MMPFC hemisphere (right) | -0.23 | 0.07 | -3.44 | **$8.360 \times 10^{-4}$** |
| Random effects | σ | | | |
| ID | 0.88 | | | |
| Residual | 0.50 | | | |
| ***Fx right FFA - amygdala*** | | | | |
| Predictors | β | SE | z | p |
| (Intercept) | 0.01 | 0.10 | 0.07 | 0.940 |
| LI | 0.07 | 0.09 | 0.78 | 0.439 |
| Amygdala hemisphere (right) | 0.01 | 0.09 | 0.08 | 0.937 |
| Age | 0.08 | 0.09 | 0.88 | 0.382 |
| Mean FD | 0.01 | 0.09 | 0.12 | 0.907 |
| Random effects | σ | | | |
| ID (σ) | 0.85 | | | |
| Residual (σ) | 0.64 | | | |
| ***Fx right FFA - STS*** | | | | |
| Predictors | β | SE | z | p |
| (Intercept) | 0.00 | 0.09 | 0.00 | 0.999 |
| LI | 0.07 | 0.07 | 0.92 | 0.358 |
| STS hemisphere (right) | -0.03 | 0.12 | -0.23 | 0.820 |
| Age | 0.06 | 0.07 | 0.79 | 0.432 |
| Mean FD | 0.17 | 0.07 | 2.30 | **0.024** |
| Random effects | σ | | | |
| ID (σ) | 0.39 | | | |
| Residual (σ) | 0.89 | | | |

*Lateralisation index was calculated at threshold p<0.10. Interaction terms are denoted with "x"; interaction terms of age-by-hemisphere-by-lateralisation index (and all embedded two-way interaction terms) for each test were not statistically significant and were excluded from the final models. Reported βs are standardised, ps<0.05 are shown in bold. N=110. FFA = Fusiform Face Area; MMPFC = Middle Medial Prefrontal Cortex; STS = Superior Temporal Sulcus; Fx = Functional Connectivity; FD = Framewise Displacement; LI = Lateralisation Index; SE = Standard Error. See Appendix S3 for further results and discussion of lateralisation index metric.*

***Table S9:*** *Associations between lateralisation of FFA response - calculated at p<0.05 - and functional connectivity between right FFA, MMPFC, amygdala and STS*

| | | | | |
|---|---|---|---|---|
| ***Fx right FFA - MMPFC*** | | | | |
| Predictors | β | SE | z | p |
| (Intercept) | 0.09 | 0.11 | 0.89 | 0.374 |
| LI | 0.11 | 0.10 | 1.05 | 0.296 |
| MMPFC hemisphere (right) | -0.03 | 0.08 | -0.41 | 0.680 |
| Age | 0.06 | 0.10 | 0.59 | 0.559 |
| Mean FD | 0.13 | 0.11 | 1.19 | 0.237 |
| Random effects | σ | | | |
| ID | 0.89 | | | |
| Residual | 0.53 | | | |
| ***Fx right FFA - amygdala*** | | | | |
| Predictors | β | SE | z | p |
| (Intercept) | -0.01 | 0.10 | -0.06 | 0.953 |
| LI | 0.03 | 0.09 | 0.32 | 0.747 |
| Amygdala hemisphere (right) | 0.06 | 0.09 | 0.63 | 0.528 |
| Age | 0.08 | 0.10 | 0.82 | 0.416 |
| Mean FD | -0.01 | 0.10 | -0.12 | 0.907 |
| Random effects | σ | | | |
| ID (σ) | 0.79 | | | |
| Residual (σ) | 0.65 | | | |
| ***Fx right FFA - STS*** | | | | |
| Predictors | β | SE | z | p |
| (Intercept) | 0.02 | 0.10 | 0.22 | 0.829 |
| LI | 0.08 | 0.07 | 1.03 | 0.305 |
| STS hemisphere (right) | -0.07 | 0.13 | -0.52 | 0.606 |
| Age | 0.02 | 0.08 | 0.22 | 0.828 |
| Mean FD | 0.15 | 0.08 | 1.93 | 0.057 |
| Random effects | σ | | | |
| ID (σ) | 0.34 | | | |
| Residual (σ) | 0.90 | | | |

*Lateralisation index was calculated at threshold p<0.05. Age-by-hemisphere-by-lateralisation index interaction terms (and embedded two-way interactions) for each test that were not statistically significant were excluded from the final models. Reported βs are standardised, ps<0.05 are shown in bold. N=97. FFA = Fusiform Face Area; MMPFC = Middle Medial Prefrontal Cortex; STS = Superior Temporal Sulcus; Fx = Functional Connectivity; LI = Lateralisation Index; FD = Framewise Displacement; SE = Standard Error. See Appendix S3 for further results and discussion of lateralisation index metric.*

***Table S10:*** *Associations between lateralisation of FFA response and functional connectivity between left FFA, MMPFC, amygdala and STS*

| ***Fx left FFA - MMPFC*** | | | | |
|---|---|---|---|---|
| Predictors | β | SE | z | p |
| (Intercept) | 0.10 | 0.10 | 1.01 | 0.316 |
| LI | 0.01 | 0.09 | 0.15 | 0.881 |
| MMPFC hemisphere (right) | -0.08 | 0.08 | -1.03 | 0.307 |
| Age | 0.12 | 0.09 | 1.26 | 0.211 |
| Mean FD | 0.07 | 0.09 | 0.80 | 0.426 |
| Random effects | σ | | | |
| ID | 0.84 | | | |
| Residual | 0.60 | | | |
| ***Fx left FFA - amygdala*** | | | | |
| Predictors | β | SE | z | p |
| (Intercept) | 0.07 | 0.10 | 0.66 | 0.510 |
| LI | 0.07 | 0.09 | 0.72 | 0.471 |
| Amygdala hemisphere (right) | -0.08 | 0.08 | -1.13 | 0.263 |
| Age | 0.02 | 0.09 | 0.19 | 0.848 |
| Mean FD | -0.07 | 0.09 | -0.73 | 0.467 |
| Random effects | σ | | | |
| ID (σ) | 0.87 | | | |
| Residual (σ) | 0.56 | | | |
| ***Fx left FFA - STS*** | | | | |
| Predictors | β | SE | z | p |
| (Intercept) | 0.11 | 0.10 | 1.14 | 0.256 |
| LI | 0.00 | 0.07 | -0.05 | 0.961 |
| STS hemisphere (right) | -0.13 | 0.13 | -0.98 | 0.328 |
| Age | -0.04 | 0.07 | -0.53 | 0.599 |
| Mean FD | 0.09 | 0.07 | 1.25 | 0.212 |
| Random effects | σ | | | |
| ID (σ) | 0.18 | | | |
| Residual (σ) | 0.99 | | | |

*Lateralisation index was calculated at threshold $p<0.10$. Interaction terms of age-by-hemisphere-by-lateralisation index (and all embedded two-way interaction terms) for each test were not statistically significant and were excluded from the final models. Reported βs are standardised. N=110. FFA = Fusiform Face Area; MMPFC = Middle Medial Prefrontal Cortex; STS = Superior Temporal Sulcus; Fx = Functional Connectivity; FD = Framewise Displacement; LI = Lateralisation Index; SE = Standard Error. See Appendix S3 for further results and discussion of lateralisation index metric.*

***Table S11:*** *Associations between lateralisation of FFA response and functional maturity of FFA, MMPFC, amygdala, and STS*

***A***

| | | | | |
|---|---|---|---|---|
| | | ***Fm MMPFC*** | | |
| Predictors | β | SE | z | p |
| (Intercept) | 0.03 | 0.09 | 0.29 | 0.771 |
| LI | -0.02 | 0.08 | -0.23 | 0.820 |
| MMPFC hemisphere (right) | -0.03 | 0.09 | -0.31 | 0.757 |
| Age | 0.36 | 0.08 | 4.54 | **$1.489 \times 10^{-5}$** |
| Mean FD | -0.18 | 0.08 | -2.28 | **0.025** |
| Random effects | σ | | | |
| ID | 0.66 | | | |
| Residual | 0.66 | | | |
| | | ***Fm amygdala*** | | |
| Predictors | B | SE | z | p |
| (Intercept) | 0.02 | 0.09 | 0.22 | 0.825 |
| LI | 0.01 | 0.09 | 0.11 | 0.909 |
| Amygdala hemisphere (right) | 0.00 | 0.09 | -0.05 | 0.958 |
| Age | 0.20 | 0.09 | 2.33 | **0.022** |
| Mean FD | 0.03 | 0.09 | 0.38 | 0.704 |
| Random effects | σ | | | |
| ID (σ) | 0.76 | | | |
| Residual (σ) | 0.64 | | | |
| | | ***Fm STS*** | | |
| Predictors | β | SE | z | p |
| (Intercept) | 0.05 | 0.09 | 0.60 | 0.551 |
| LI | -0.09 | 0.07 | -1.33 | 0.185 |
| STS hemisphere (right) | -0.05 | 0.12 | -0.44 | 0.659 |
| Age | 0.22 | 0.07 | 3.33 | **0.001** |
| Mean FD | -0.07 | 0.07 | -1.05 | 0.294 |
| Random effects | σ | | | |
| ID (σ) | 0.22 | | | |
| Residual (σ) | 0.92 | | | |

***B***

| | | | | |
|---|---|---|---|---|
| | | ***Fm FFA*** | | |
| Predictors | β | SE | z | p |
| (Intercept) | 0.02 | 0.09 | 0.26 | 0.795 |
| LI | 0.02 | 0.07 | 0.34 | 0.734 |
| MMPFC hemisphere (right) | 0.00 | 0.11 | -0.01 | 0.993 |
| Age | 0.30 | 0.07 | 4.22 | **$5.086 \times 10^{-5}$** |
| Mean FD | -0.17 | 0.07 | -2.49 | **0.014** |
| Random effects | σ | | | |
| ID | 0.41 | | | |
| Residual | 0.84 | | | |

*Lateralisation index was calculated at threshold p<0.10. Age-by-hemisphere-by-lateralisation index interaction terms (and embedded two-way interactions) for each test were not statistically significant and excluded from the final models. Reported βs are standardised, ps<0.05 are shown in bold. N=110. FFA = Fusiform Face Area; MMPFC = Middle Medial Prefrontal Cortex; STS = Superior Temporal Sulcus; Fm = Functional Maturity; LI = Lateralisation Index; FD = Framewise Displacement; SE = Standard Error. See Appendix S3 for further results and discussion of lateralisation index metric.*

***Figure S1:*** *Significant associations between age and response magnitude of fROIs to face and scene events*

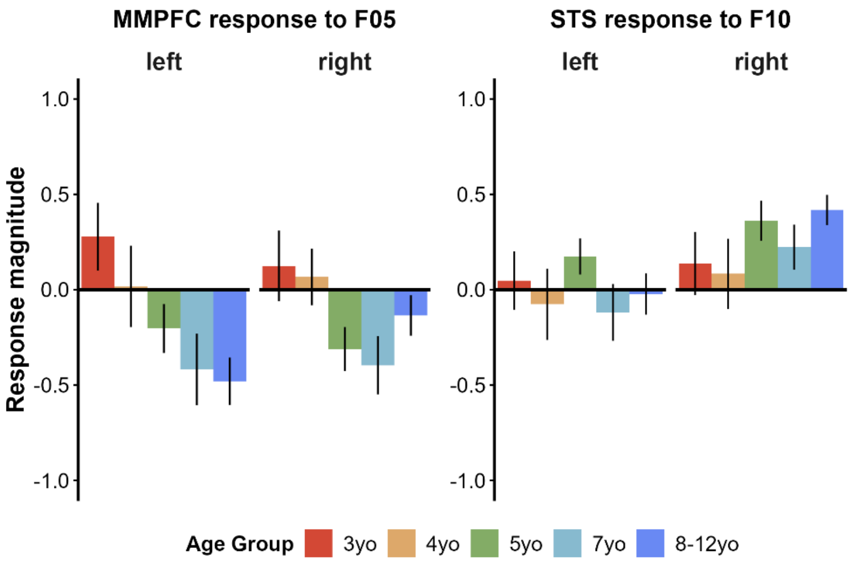


*Response magnitude (y-axis) in both left and right MMPFC decreased with age during a face scene where Peck's head is covered in porcupine spines and Gus removes them (F05). Response magnitude in both left and right STS increased with age during a face scene where Gus scolds the crocodile (F10). These findings are consistent with previous analyses of this dataset: MMPFC responses to non-preferred categories (i.e., events depicting others' physical pain/bodily sensations) decrease with age (Richardson et al., 2018), while STS shows age-related increases in response to adult-defined face events (Kamps, Richardson et al., 2022). fROIs = Functional Regions Of Interest.*

***Figure S2:*** *Inter-region correlations (i.e., functional connectivity)*

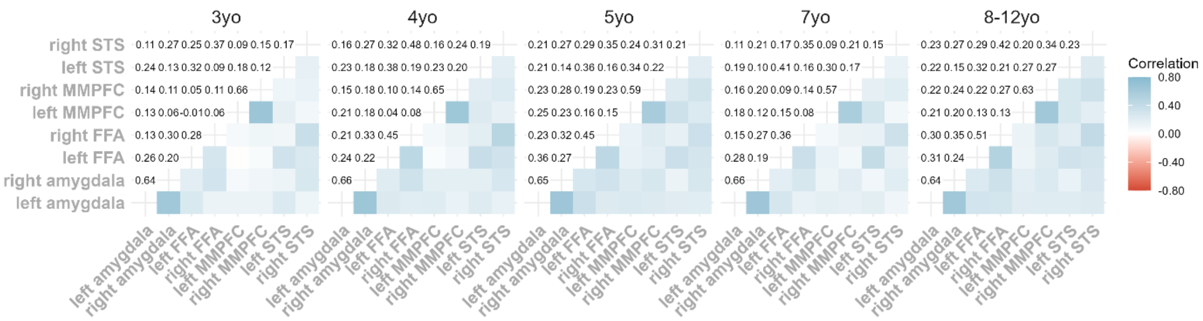


*Average z-scored Pearson's correlation matrices across all regions of interest per age group (3yo: n=15; 4yo: n=14; 5yo: n=32; 7yo: n=23; 8-12yo: n=33). Connectivity between right FFA and right MMPFC increased more with age than that between right FFA and left MMPFC. In addition, connectivity between the right MMPFC and bilateral STS increased with age. This finding is consistent with previous analyses of this dataset suggesting that STS development reflects an increasing integration with other higher-level social cortical regions with age (Kamps, Richardson et al., 2022).*

***Figure S3:*** *Age and functional connectivity between left FFA, MMPFC, amygdala and STS*

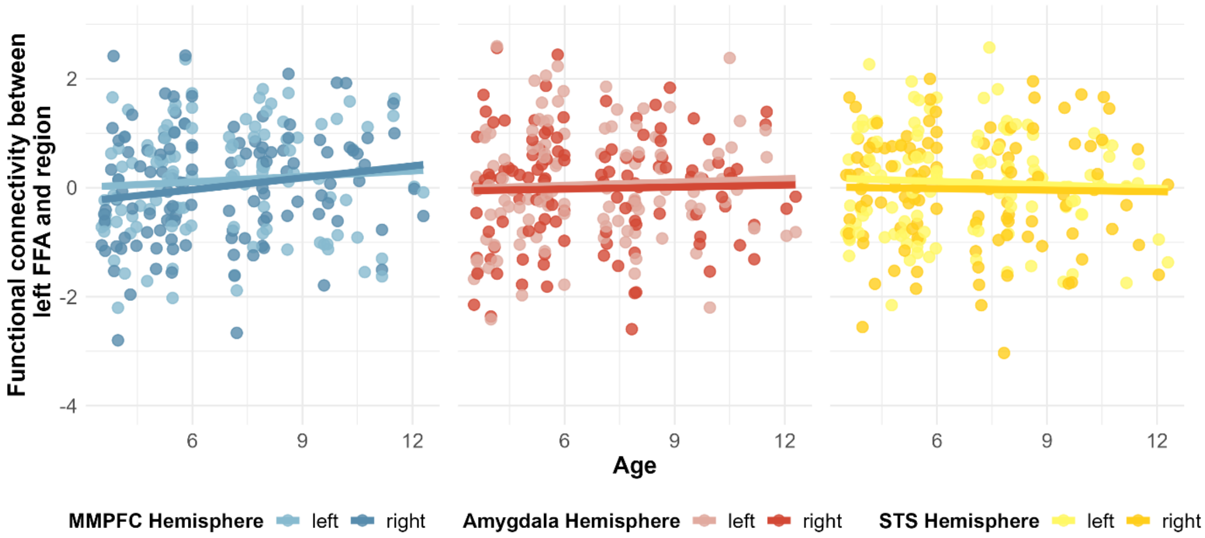


*Association between age (x axis) and functional connectivity between left FFA and MMPFC (blue), amygdala (red), or STS (yellow; y axis). In all scatterplots, lighter colours correspond to the left hemisphere, darker colours correspond to the right hemisphere. Lines represent linear regression fits estimated using the least-squares method. N=117. FFA = Fusiform Face Area; MMPFC = Middle Medial Prefrontal Cortex; STS = Superior Temporal Sulcus.*

***Figure S4:*** *Functional maturity of left FFA and functional maturity of/connectivity with MMPFC, amygdala and STS*

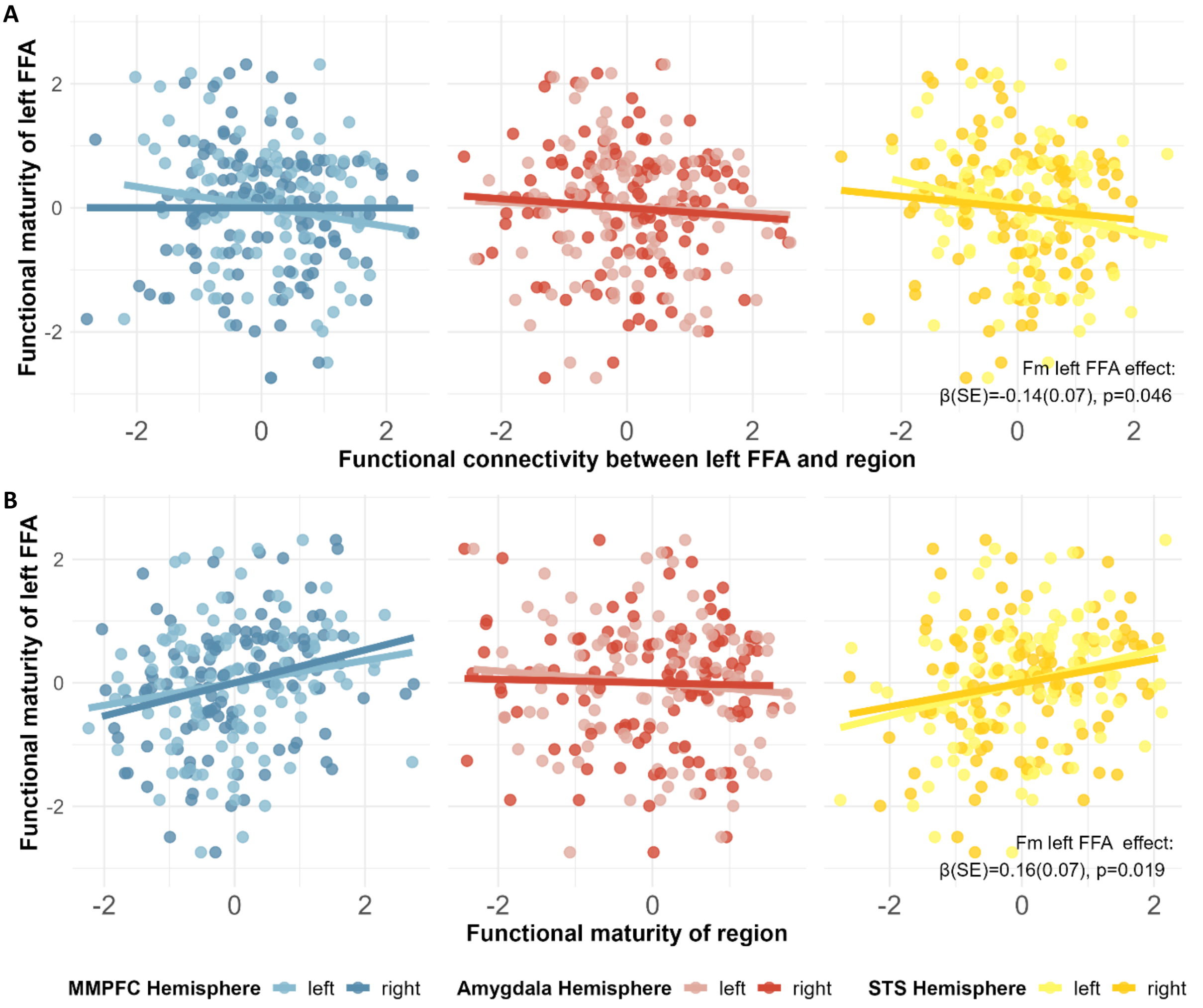


*A) Associations between functional connectivity (x axis) between left FFA and MMPFC (blue), amygdala (red), and STS (yellow) and functional maturity of left FFA (y axis). B) Associations between functional maturity of MMPFC, amygdala and STS (x axis) and functional maturity of left FFA (y axis). In all scatterplots, lighter colours correspond to the left hemisphere, darker colours correspond to the right hemisphere of MMPFC, amygdala and STS. Lines represent linear regression fits estimated using the least-squares method. N=117. FFA = Fusiform Face Area; MMPFC = Middle Medial Prefrontal Cortex; STS = Superior Temporal Sulcus; Fm = Functional Maturity; SE = Standard Error.*

***Figure S5:*** *Correlations of functional connectivity between FFA, MMPFC, amygdala and STS*

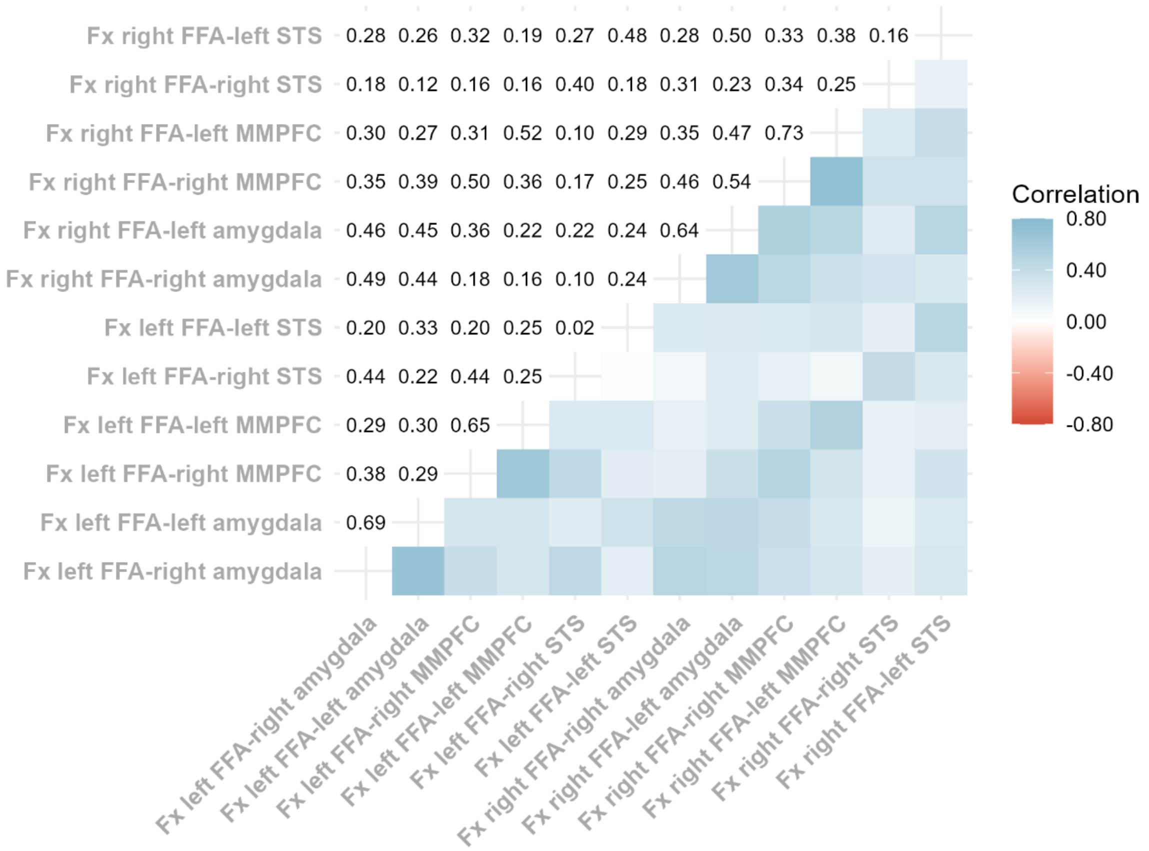


*Average correlation matrix (Pearson's r) across all functional connectivity measures. N=117. Fx = Functional Connectivity.*

***Figure S6:*** *Correlations of functional maturity between FFA, MMPFC, amygdala and STS*

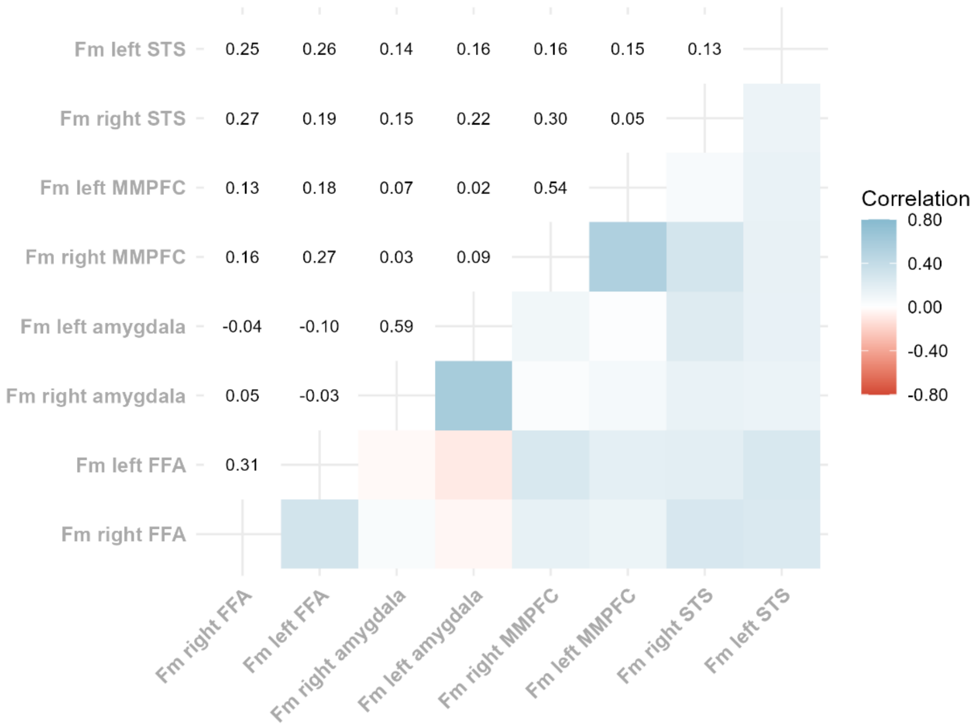


*Average correlation matrix (Pearson's r) across all functional maturity measures. N=117. Fm = Functional Maturity.*

***Figure S7:*** *Age and FFA response lateralisation*

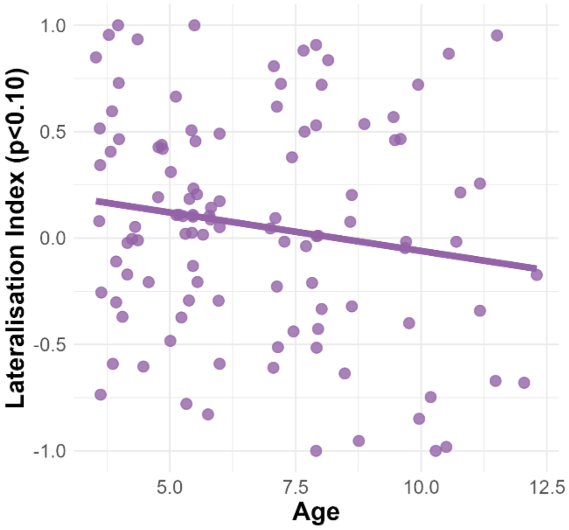


*Scatterplot shows lateralisation index of FFA, calculated at p<0.10 (y axis) by age (in years, x axis) in 3- to 12-year-old children. Lines represent linear regression fits, estimated using the least-squares method. N=110. See Appendix S3 for further results and discussion of lateralisation index metric.*

***Figure S8:*** *Lateralisation of FFA response and functional connectivity of right FFA to/maturity of MMPFC, amygdala, and STS*

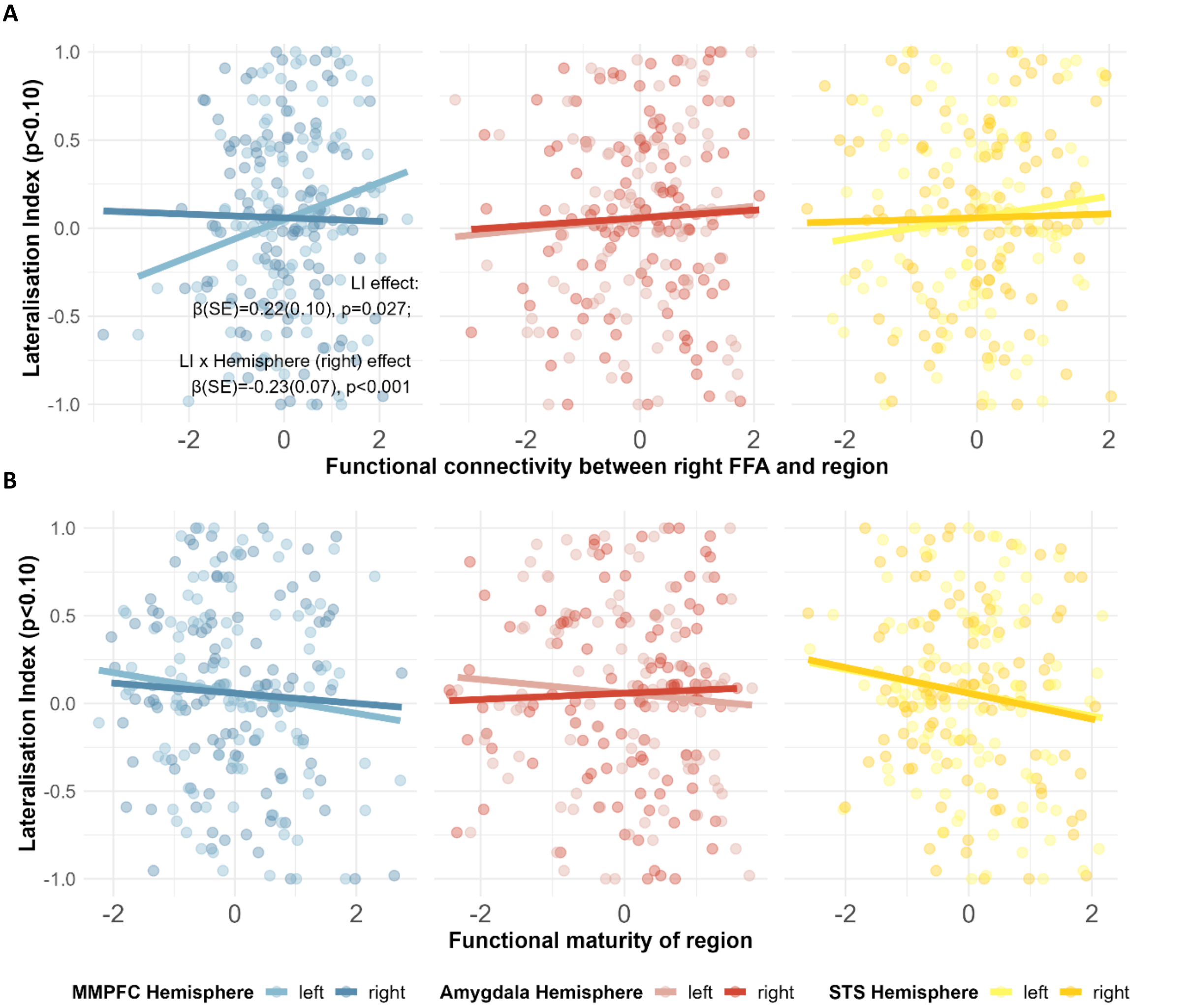


*Associations between functional connectivity (x axis) between left FFA and MMPFC (blue), amygdala (red) and STS (yellow) and lateralisation index, calculated at threshold p<0.10 (y axis). In all scatterplots, lighter colours correspond to the left hemisphere, darker colours correspond to the right hemisphere of MMPFC, amygdala and STS. Lines represent linear regression fits estimated using the least-squares method. N=110. FFA = Fusiform Face Area; MMPFC = Middle Medial Prefrontal Cortex; STS = Superior Temporal Sulcus; LI = Lateralisation Index; SE = Standard Error. See Appendix S3 for further results and discussion of lateralisation index metric.*

***Figure S9:*** *Lateralisation of FFA response - calculated at p<0.05 - and functional connectivity of right FFA to MMPFC, amygdala and STS*

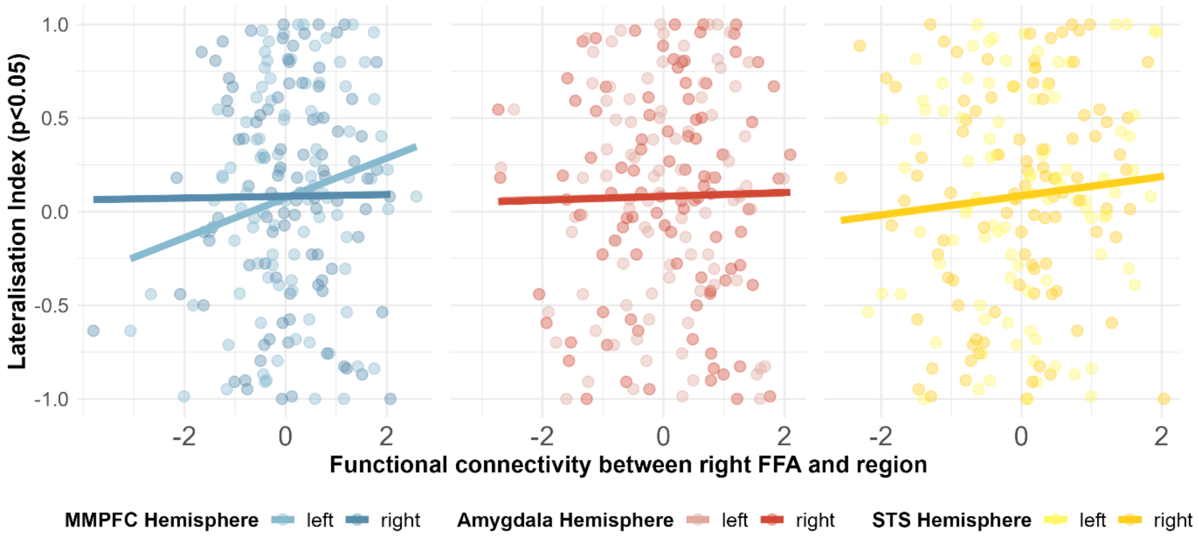


*Associations between functional connectivity (x axis) between left FFA and MMPFC (blue), amygdala (red) and STS (yellow) and lateralisation index, calculated at threshold p<0.05 (y axis). In all scatterplots, lighter colours correspond to the left hemisphere, darker colours correspond to the right hemisphere of MMPFC, amygdala and STS. Lines represent linear regression fits estimated using the least-squares method. N=97. FFA = Fusiform Face Area; MMPFC = Middle Medial Prefrontal Cortex; STS = Superior Temporal Sulcus. See Appendix S3 for further results and discussion of lateralisation index metric.*

***Figure S10:*** *Lateralisation of FFA response and functional connectivity of left FFA to MMPFC, amygdala and STS*

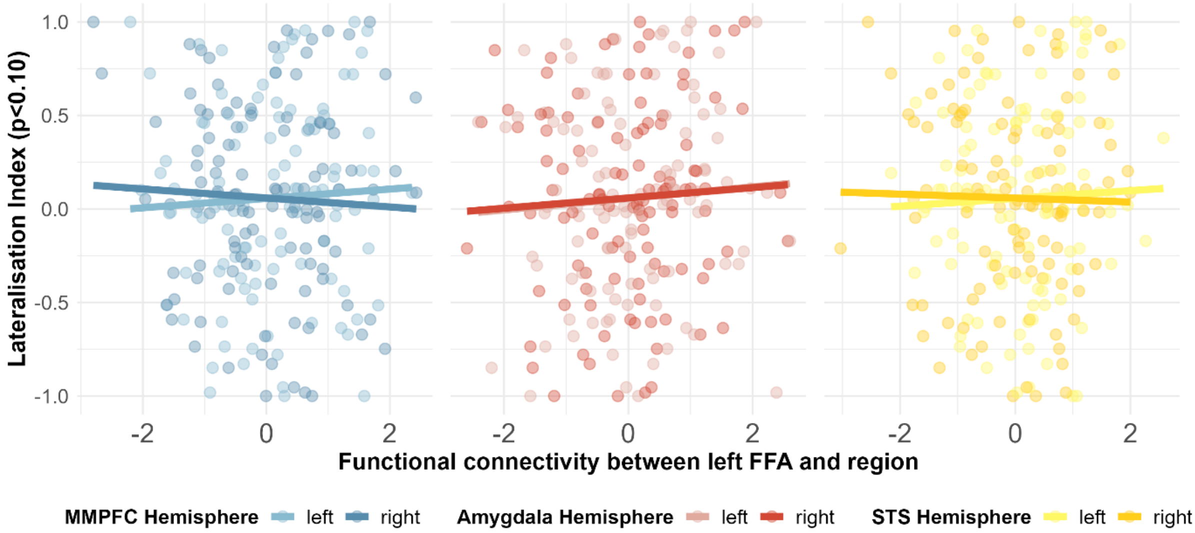


*Associations between functional connectivity (x axis) between left FFA and MMPFC (blue), amygdala (red) and STS (yellow) and lateralisation index, calculated at threshold p<0.10 (y axis). In all scatterplots, lighter colours correspond to the left hemisphere, darker colours correspond to the right hemisphere of MMPFC, amygdala and STS. Lines represent linear regression fits estimated using the least-squares method. N=110. FFA = Fusiform Face Area; MMPFC = Middle Medial Prefrontal Cortex; STS = Superior Temporal Sulcus. See Appendix S3 for further results and discussion of lateralisation index metric.*
